# IL-22 promotes genesis of small intestinal secretory cells that protect against cholera in mice

**DOI:** 10.1101/2025.09.12.675873

**Authors:** Masataka Suzuki, Yuko Hasegawa, Hailong Zhang, Zhu Liang, Guodong Tie, Ramesh A. Shivdasani, Matthew K. Waldor

**Author notes:** Department of Molecular Biology and Genetics, Cornell University, Ithaca, NY, 14853. Department of Pharmacology and Immunology, Medical University of South Carolina, Charleston, SC, 29425, USA.

## Abstract

The diarrheal disease cholera remains a global threat, but there is limited knowledge of the innate immune defenses in the small intestine that protect against the causative agent, *Vibrio cholerae*. Here, single-cell RNA-sequencing of epithelial and immune cells mapped gene expression patterns in the infant mouse small intestine and revealed changes in response to *V. cholerae* infection and prophylactic treatment with an IL22 Fc-fusion protein. Infection increased the abundance of an enterocyte subtype with high expression of defense-associated functions and stimulated production of IL22, a cytokine linked to epithelial integrity, from group 3 innate lymphoid cells. Administration of IL22Fc increased production of vibriocidal Reg3β from enterocytes and the abundance of secretory lineage and Muc2-producing goblet cells, which secreted mucus into the intestinal crypts, impairing *V. cholerae* association with the epithelium. These IL22-mediated responses limited *V. cholerae* intestinal colonization and protected mice from diarrhea and death. Our findings suggest enterocyte specialization in mucosal defense.

**Summary:** Single cell studies uncovered specialization of epithelial cells in intestinal defense. The epithelial cell-targeting cytokine IL22 protected mice from cholera by promoting the genesis of mucus secreting cells.

## Introduction

*Vibrio cholerae*, the cholera pathogen, infects several million people and causes tens of thousands of deaths annually, particularly in low- and middle-income countries in south Asia and Africa^1^. Cholera patients can lose many liters of diarrheal fluid within hours and die from dehydration without proper treatment^2^. *V. cholerae* is a curved, highly motile Gram-negative rod that associates with the epithelium of the human small intestine (SI), the site of colonization. Various *V. cholerae* genes and processes, including its Tcp pilus along with flagellar-based motility are important for intestinal colonization^3^. While replicating along the epithelium, the pathogen secretes cholera toxin, whose activities largely account for the secretory diarrhea characteristic of cholera.

Knowledge of adaptive immune responses to natural and experimental *V. cholerae* infection in humans has expanded in recent years^4^. Infection leads to the generation of protective antibody responses that can limit intestinal colonization and diarrhea^5^. However, despite more than a century of studies of cholera pathogenesis^6–8^, the comprehensive understanding of epithelial and innate immune responses to *V. cholerae* infection in the small intestine is fairly limited, though human biopsy studies have suggested roles for neutrophil infiltration, and elevated abundance of inflammation-associated and antimicrobial proteins including TNF, IL1β, IL23, lactoferrin, LPLUNC1 and defensin^6–10^. Epithelial cells represent a critical barrier preventing pathogens or commensal bacteria from entering host tissues, and these cells produce physical (e.g., mucus) and chemical (e.g., reactive oxygen species, antimicrobial peptides) obstacles that impede microbial invasion of deeper tissues^6^; epithelial cells also initiate immune responses when the epithelial barrier is compromised^11^. In cholera naïve individuals, such innate responses are likely to be critical determinants of survival given that severe cholera can kill infected individuals within a day of infection, before adaptive immune responses are generated^2^.

Modeling human pathogenesis, *V. cholerae* robustly colonizes the small intestine (SI) of infant mice without antibiotic pre-treatment. Furthermore, colonization of the infant mouse SI depends on the Tcp pathogenicity island, as is the case in humans^12^. Here, we characterized intestinal epithelial and immune cell responses to *V. cholerae* infection in infant mice using single-cell RNA sequencing (scRNA-seq). Our findings uncover patterns of mucosal innate defense and reveal that IL22 therapy shifts stem cell differentiation of intestinal stem cell daughters toward the secretory lineage, which protects against infection by impairing pathogen association with the epithelium. These findings could point to host-directed approaches for cholera treatment.

## Results

### *V. cholerae* infection triggers antimicrobial responses in intestinal epithelial cells

scRNA-seq analysis of epithelial cells harvested from the SI of postnatal day 5 (P5) infant mice revealed 15 cell clusters, including various enterocyte subtypes, goblet cells, and enteroendocrine cells (Fig. 1ab, Extended Data Fig. 1ab, Supplementary Table 1). The expression profiles in the 11 enterocyte clusters and stem cells likely distinguish where they are located along the crypt-villus axis^13^ (Extended Data Fig. 1c). However, Paneth and tuft cells were not detected (Extended Data Fig. 1d), revealing the developmental immaturity of the epithelial defense system in P5 mice^13^.

*V. cholerae* infection altered the transcriptional profiles in nearly all clusters of SI epithelial cells, particularly in the distal SI, where the pathogen burden is 10-100x greater than in the proximal SI^14^ (Extended Data Fig. 2a, Supplementary Table 2, 3). Pathway analyses showed prominent upregulation of inflammatory signaling and hypoxia pathways across almost all epithelial cell clusters (Fig. 1cd, Extended Data Fig. 2b). There were also cell type-specific infection-induced perturbations, such as in metabolic pathways (Fig. 1d, Extended Data Fig. 2c), suggesting that infection stimulates distinct regulatory networks across epithelial cell types; for example, stem cells exhibited prominent increases in expression of genes linked to the MYC pathway, suggesting that infection triggers stem cell proliferation (Extended Data Fig. 2d). Highly induced genes in stem cells were generally different than those in most enterocyte populations and goblet cells (Fig. 1c, Extended Data Fig. 2e).

**Figure 1.**
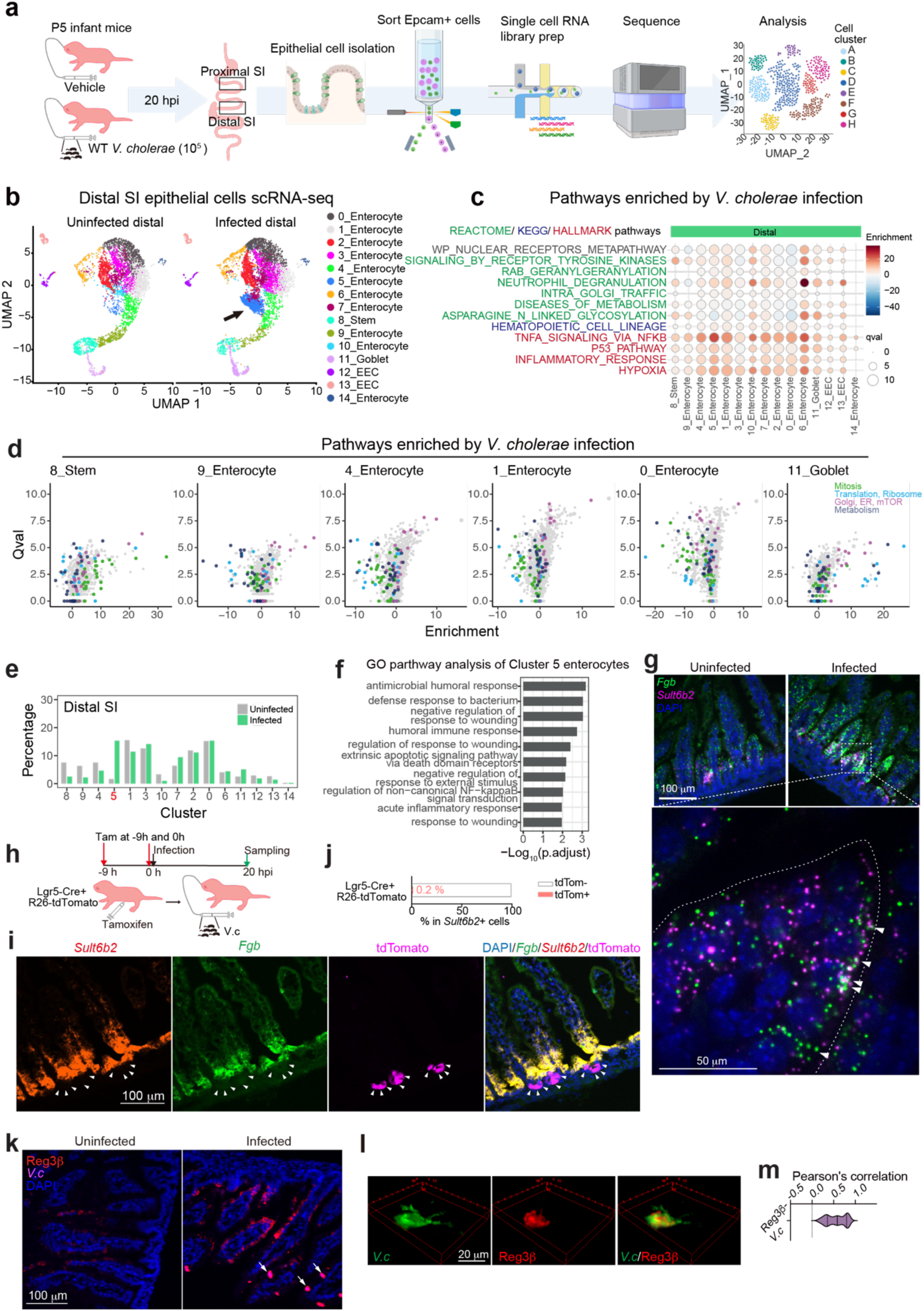
***V. cholerae* infection increases the abundance of cluster 5 enterocytes in the small intestine and elicits antimicrobial responses in intestinal epithelial cells in infant mice. (a)** Experimental scheme of scRNA-seq analysis of epithelial cells. Infant mice (P5) (n=3/group) were infected with *V. cholerae* and sacrificed 20 hours post-infection (hpi). Single-cell RNA libraries were prepared from sorted Epcam+ cells isolated from the proximal and distal small intestine (SI). **(b)** UMAP of the distal small intestine epithelial single-cell RNA-seq data from uninfected and *V. cholerae* infected P5 infant mice (n=3 for each condition, cells were pooled). EEC (enteroendocrine cells). The arrow indicates cluster 5 enterocytes in infected animals in the distal SI. (**c)** Top-hit pathways from the Single Cell Pathway Analysis (Qval > 8.75 in any cluster). Dot plot represents enrichment in color and Qval in circle size. Stem (stem cells); EC (enterocytes); Gob (goblet cells); EEC (enteroendocrine cells). (**d)** Volcano plot showing q-value (Qval) and pathway enrichment from Single Cell Pathway Analysis. Pathway enrichment indicates a mean pathway change comparing uninfected and *V. cholerae*-infected conditions. Colors represent the following pathways: Mitosis (green), Translation and Ribosome (light blue), Golgi, ER, and mTOR (pink), and Metabolism (dark blue). (**e)** Percentage of cells in each cluster vs the total cell number in the distal SI. (**f)** GO pathway analysis of cluster 5 representative genes. Enrichment analysis was performed using gost function in gProfiler2, based on the hypergeometric test with Benjamini–Hochberg false discovery rate (FDR) correction for multiple testing. **(g)** *Fgb* and *Sult6b2* gene detection by single molecule RNA FISH co-stained with DAPI (blue) in the distal SI tissue at 20 hpi. White arrowheads in the higher magnification image indicate the expression of *Fgb* (green) and *Sult6b2* (magenta) mRNA in epithelial cells. Dotted line indicates the epithelial cell membrane. Signal intensity was adjusted to visualize single puncta in the high magnification image. Color-separated images are shown in Extended Data Fig.3b. Shown are representative images of 29 images from 4 uninfected animals and 34 images from 4 infected animals. (**h**) Experimental protocol for use of lineage tracing to study the origins of cluster 5 cells. To trace the Lgr5 lineage with a pulse of tdTomato labeling, P5 infant mice were treated with 2 doses of tamoxifen (Tam) at −9 and 0 hpi and the small intestine was collected at 20 hpi. (**i**) RNA FISH for cluster 5 marker genes *Sult6b2* (red) and *Fgb* (green), protein staining for tdTomato (magenta), and DAPI (blue). White arrowheads show tdTomato+ cells. Representative images of 160 images from 4 animals are shown. (**j**) Fraction of tdTomato positive cells (pink) and tdTomato negative cells (white) among *Sult6b2*+ cells. 160 images from 4 animals were analyzed. Data are presented as mean ± SD. (**k)** Immunostaining of Reg3β (red), *V. cholerae* (magenta), and DAPI (blue) in the distal SI from an uninfected animal (left) or *V. cholerae*-infected animal (right). Shown are representative images of 66 images from 3 uninfected animals and images of 47 images from 3 infected animals. Scale bar: 100 μm. White arrows show the colocalization of Reg3β and *V. cholerae* on the epithelium. (**l)** Deconvoluted z-stack image of *V. cholerae* (green) and Reg3β (red) from a puncta of Reg3β/*V. cholerae* colocalization image. Overlay image was taken by confocal microscope using ×100 lens and z-stack with 0.3 micron gaps per layer. Representative images from 7 images are shown. **(m)** Pearson’s correlation coefficient between Reg3β and *V. cholerae* co-localization using the images of (**k**) from infected animals. 55 images from 4 animals were analyzed and plotted in a violin plot. (Fig. 1a is partially created in BioRender. Waldor, M. (2026) https://BioRender.com/shf91td)

There were minimal differences in the relative abundances of epithelial clusters in the proximal SI in control vs infected animals (Extended Data Fig. 1b, 2f). In contrast, there was a marked increase in the abundance (12.6-fold) of cluster 5 enterocytes in the distal SI of infected animals (Fig. 1b, e). Cluster 5 enterocytes expressed genes encoding Reg3β and Reg3γ antimicrobial peptides even in the absence of infection (Extended Data Fig 3a), suggesting that this epithelial subset is specialized for defense against infection. With infection, cluster 5 enterocyte highly expressed genes classed as defense responses, including *Il18*, as well as genes encoding fibrinogen proteins, *Fga* and *Fgb*, which are traditionally associated with clotting^15^ (Fig. 1f, Extended Data Fig 3a). In contrast, infection was associated with reduced expression of several metabolic pathways in this enterocyte subset (Extended Data Fig. 2c). Single molecule RNA FISH using probes for cluster 5 marker genes confirmed that these cells were found in the epithelium and that their abundance increased with infection; cluster 5 enterocytes were primarily located toward the base of the villus (Fig 1g and Extended Data Fig 3b), the site where *V. cholerae* primarily localizes as well^14^.

Since the abundance of cluster 5 enterocytes increased markedly, we conducted lineage tracing to investigate whether the infection-associated increase in abundance of cluster 5 enterocytes arose from Lgr5+ stem cells during the 20-hour infection period or from enterocytes already present on villi. Lgr5^Cre(ER-T2)^/ Rosa26R^LSL-TdTomato^ animals, which express a Lgr5-dependent tamoxifen-inducible tdTomato, were infected with *V. cholerae*. After 20 hours of infection, only 0.2 % of cluster 5 (*Sult6b2*+) cells were tdTomato+ (Fig. 1h-j). To test whether cluster 5 cells were derived from transient amplifying cells or had themselves expanded, we administered the S-phase reagent EdU at the time of infection or 2 hours before tissue harvest. When tissues were examined, only 5.2 % or 0.8 % of cluster 5 (*Sult6b2*+) cells were EdU+ when this reagent was administered at the time of infection or 2 hours prior to tissue sampling respectively (Extended Data Fig. 3c-e). Together these findings indicate that the infection-associated expansion of cluster 5 enterocytes originates from another enterocyte population and did not differentiate from stem cells, or self-amplify in response to the infection. Moreover, RNA velocity analysis of our scRNA-seq data corroborated this idea, predicting that cluster 5 cells originated from *Lgr5*^-^negative, *MKi67*-negative cells (Extended Data Fig. 3f), and buttressing the conclusion that the majority of cluster 5 cells originate from another subset of enterocytes.

Infection stimulated increased expression of *Reg3b* and *Reg3g* in all the epithelial cell clusters in the distal SI (Extended Data Fig 3a). Immunostaining established that Reg3β is highly expressed in epithelial cells towards the upper crypt and middle section of villi in infected animals (Fig. 1k, Extended Data Fig 3g).

Moreover, Reg3β staining was often colocalized with *V. cholerae* microcolonies found on the epithelial surface (Fig.1k-m, Extended Data Fig 3g), suggesting that this lectin is secreted and bound to the pathogen during infection. Together, these analyses reveal that *V. cholerae* infection can shifts the functions of enterocytes towards defense both by reprogramming gene expression in many enterocyte populations and by increasing the abundance of a distinct, defense-associated subset (cluster 5).

### *V. cholerae* infection stimulates IL22 expression in type 3 innate lymphoid cells

Crosstalk between epithelial and immune cells found in the SI lamina propria plays a key role in host protection against enteric bacterial infection^11,16^. We also carried out scRNA-seq analysis of CD45+ immune cells isolated from the SI lamina propria of control and *V. cholerae*-infected P5 infant mice. Cell cluster analysis based on marker gene expression revealed 32 clusters in both the proximal and distal SI (Fig. 2a, Extended Data Fig. 4a-c, Supplementary Table 1). The cell types previously observed in adult animals^17^ were also detectable in the SI of P5 neonates with the exception of plasma cells (Extended Data Fig. 4d). As observed for epithelial cell responses, there were more differentially expressed genes in the CD45+ cells isolated from the distal vs. proximal SI and considerable variability in infection-induced changes in gene expression among cell clusters (Fig. 2b, Extended Data Fig. 5a, Supplementary Table 4, 5). Pathway analysis revealed that, in contrast to scRNA-seq studies in *V. cholerae*-infected zebra fish, where the pathogen reduced interferon signaling in the intestine^18^, interferon response pathway genes were prominently induced in B and T cells, which express the receptors for Type I (α and β) and II (γ) interferon, *Ifnar1/Ifnar2* and *Ifngr1/Ifngr2* respectively, while innate lymphoid cells responded to pro-inflammatory IL17, IL23 signaling but not to interferons (Fig. 2b-d, Extended Data Fig. 5bc). NK cells may be the source of interferon gamma (Extended Data Fig. 5c), but producer(s) of Type I and III (λ) interferon were not apparent.

**Figure 2.**
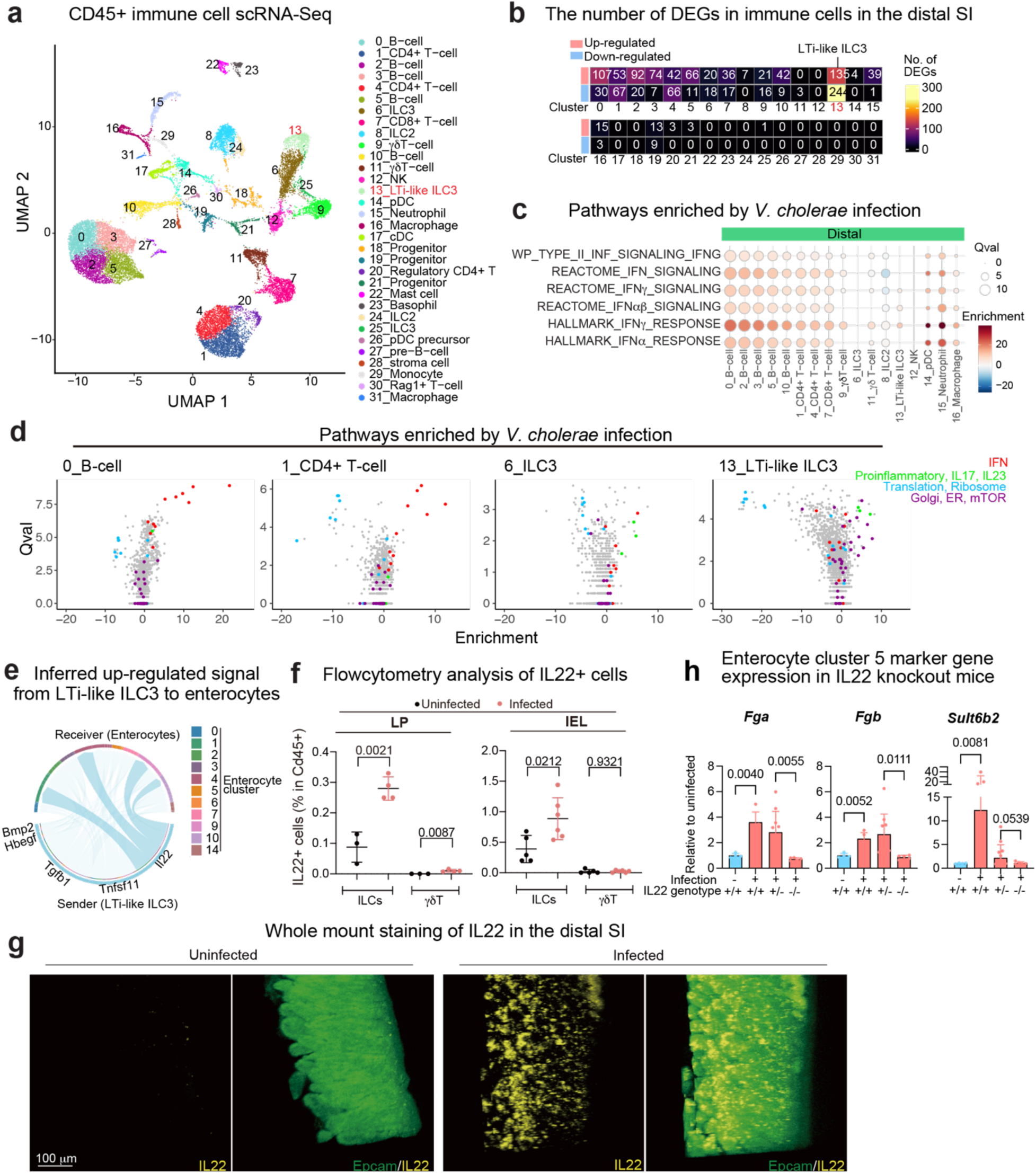
***V. cholerae* infection induces IL22 expression in Type 3 innate lymphoid cells. (a)** UMAP of CD45+ immune cells (23,836 cells including all conditions: proximal and distal SI, mock and infected samples) in the lamina propria of the small intestine. **(b)** Heatmap of the numbers of differentially expressed genes (absolute Fold Change > 1, adjusted p-value < 0.00001, genes on chromosome X and Y were excluded as both, male and female infant mice were used for each condition, in each cluster). (**c)** Dot plots showing enrichment and Qval of the Single Cell Pathway Analysis results in color and circle size, respectively. Clusters with more than 500 cells (total cell number) isolated from distal SI are shown. **(d**) Volcano plots showing enrichment and Qval of Single Cell Pathway Analysis results in the representative clusters for B cells, T cells, and ILC3s. Colors represent pathways “Interferon” for IFN (red), Proinflammatory, IL17, and IL23 (green), Translation and Ribosome (blue), Golgi, ER, and mTOR (purple). **(e)** CellChat^19^-based analysis of up-regulated signaling pathways from LTi-like ILC3 to enterocytes. IL22-IL22Ra1/IL10Rb signaling is highlighted. Color code represents the types of enterocytes in the receiver. **(f)** IL22+ cell populations in CD45+ cells in lamina propria (LP, uninfected: n = 3, infected: n = 4) and intraepithelial lymphocyte (IEL, uninfected: n = 5, infected: n = 6). Data are represented as mean ± SD for biological replicates. Two-sided unpaired t-test. Data are representative of two independent experiments. **(g)** Whole mount staining of IL22 (yellow) and Epcam (green) in uninfected or infected distal SI. Images are representative of three independent samples/group. (**h**) qPCR analysis of cluster 5 enterocyte marker gene (*Fga*, *Fgb*, and *Sult6b2*) expression in the distal SI from uninfected WT C57BL/6 (*IL22^+/+^*) (n = 4), *V. cholerae* infected WT C57BL/6 (*IL22^+/+^*) (n = 5), infected *IL22^+/-^* (n = 10) and infected *IL22^-/-^* (n = 5) littermate animals. Expression levels were normalized to β-Actin expression and shown as relative expression to uninfected *IL22^+/+^* group. Data are represented as mean ± SD. Two-sided Brown-Forsythe and Welch ANOVA.

Differential expressed gene (DEG) analyses uncovered that gene expression patterns in Type 3 LTi-like innate lymphoid cell (LTi-like ILC3) are prominently altered by infection (Fig. 2b). Cell-cell communication inference^19^ was used to identify potential signaling between lamina propria LTi-like ILC3 and epithelial cells. Several pathways were inferred, including IL22-IL22ra1/IL10rb signaling from LTi-like ILC3 to epithelial cells (Fig. 2e). We focused additional analyses on IL22 signaling because this cytokine was highly up-regulated by *V. cholerae* infection (> 16-fold increase, adjusted p-value 3.24×10^-38^) (Supplementary Table. 5) and is known to promote epithelial barrier integrity^20–24^. Flow cytometry analysis and whole mount staining corroborated the increased number of IL22+ cells, which were primarily ILCs and not γ8T cells and found in the lamina propria and among intraepithelial lymphocytes (IEL) (Fig. 2fg, Extended Data Fig. 5d). Quantitative PCR analysis with IL22 knockout animals confirmed that *V. cholerae* infection-induced expression of *Reg3b* and *Reg3g* depends on IL22 signaling (Extended Data Fig. 5e), as observed in studies with other enteric pathogens ^21,25^. Furthermore, cluster 5 enterocyte marker genes induced by *V. cholerae* infection (Fig. 1g, Extended Data Fig. 3a) were not induced in IL22 knockout mice (Fig. 2h), suggesting that IL22 signaling promotes the generation of this defense-associated enterocyte subset.

Thus, similar to epithelial cell responses, transcriptional responses to *V. cholerae* infection in lamina propria immune cells were cell type-specific and more pronounced in the distal SI. Also, among lamina propria innate immune cells, increased ILC3 expression of regulatory cytokines, including IL22, was notable.

### Il22 protects infant mice from *V. cholerae* intestinal colonization and diarrhea

Since IL22 has been linked to intestinal defense^20–24^ and transcriptional profiling suggested that *V. cholerae* infection stimulates an IL22-based signaling axis between lamina propria ILC3 cells and the epithelium, we investigated whether this cytokine could impact the course of *V. cholerae* infection in suckling mice. Because of the low expression of IL22 in very young animals^26^, we speculated that the administration of exogenous IL22 could change the outcome of *V. cholerae* infection in P5 animals. Infant animals were given two doses of an IL22Fc protein^27^ at 24 and 0 hours before challenge with *V. cholerae* (Fig. 3a). This prophylactic regimen of IL22Fc administration markedly decreased the *V. cholerae* burden in both the proximal and distal SI (Fig 3b, Extended Data Fig. 6ab). In contrast, administration of recombinant Il17a or Il17f, which were also activated in LTi-ILC3 by *V. cholerae* infection, or Ifnβ or Ifnγ, whose receptor downstream signaling was activated in T, B and myeloid cells (Fig. 2cd, Extended Data Fig. 5b), did not influence *V. cholerae* colonization (Extended Data Fig. 6a-d).

**Figure 3.**
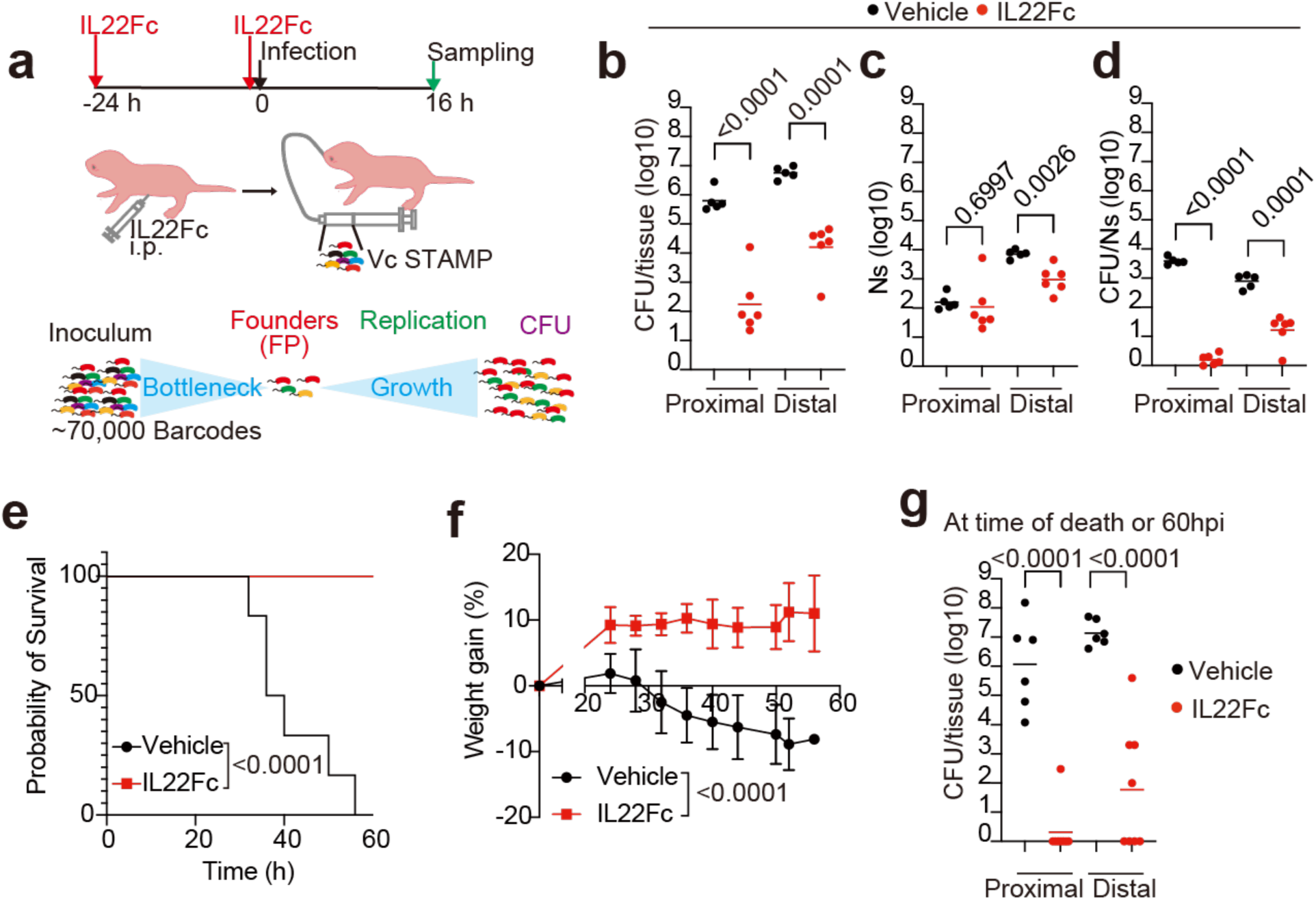
IL22 treatment protects infant mice from experimental cholera. **(a)** Schematic of IL22 prophylactic regime. A barcoded library of *V. cholerae* was used in these experiments to enable measurement of the founding population (Ns), the number of cells from the inoculum that establish infection, and CFU/Ns which represents the net replication of the founders during infection. **(b-d)** CFU (b), founding population (Ns) (c), and replication (CFU/Ns) (d) of *V. cholerae* in the proximal and distal SI. Vehicle: PBS-treated animals, IL22Fc: IL22Fc-treated animals. Bars indicate the geometric mean for biological replicates (Vehicle: n = 5, IL22Fc: n = 6). Two-sided unpaired t-test after log transformation. (**e)** Kaplan-Meyer plot of infant mice survival in *V. cholerae* infection in IL22Fc-treated or control animals; vehicle control (black, n = 6), IL22Fc-treated group (red, n = 8). Log-rank test. (**f)** Percent weight gain relative to the time of infection; vehicle (black, n = 6), IL22Fc-treated group (red, n = 8). Data are represented as mean ± SD. Two-way ANOVA. **(g)** *V. cholerae* burden in the proximal and distal SI at the time of death (vehicle-treated, black, n = 6) or 60 hpi (IL22Fc-treated, red, n = 8). Bars indicate the geometric mean. Two-sided unpaired t-test. P-values are shown.

A bacterial-barcoding strategy was used to determine whether IL22-treatment strengthens the initial bottleneck that limits *V. cholerae* colonization and/or prevents subsequent replication. Amplicon sequencing and the STAMPR analytic pipeline monitor the diversity of otherwise isogenic barcoded bacteria to quantify the number of cells that survive colonization bottlenecks to initiate infection, referred to as the ‘founding population’^28^. STAMPR revealed that both the founding population (Ns) and net replication (CFU/Ns) in the distal SI were reduced in treated animals (Fig. 3cd). These results suggest that IL22Fc both limits net replication and tightens the host bottleneck, restricting the number of niches in the distal SI that are permissive for *V. cholerae* replication or killing cells before they become founders.

When *V. cholerae*-infected animals are left with a dam and monitored for survival, they develop diarrhea, lose weight, and generally die within 60 hours (Fig. 3e-g, Extended Data Fig. 6e). In marked contrast, in animals treated with IL22Fc, no weight loss, diarrhea, or death was observed (Fig. 3e-g). In fact, the treated mice gained weight and at 60 hours had either entirely cleared *V. cholerae* or had dramatically lower pathogen burdens than observed in the moribund control animals (Fig. 3h). Only a single dose of IL22Fc 24 or 48 hours before infection was sufficient to impair *V. cholerae* colonization (Extended Data Fig. 6ab), while IL22Fc treatment 5 days prior to infection had limited effect (Extended Data Fig. 6f). Administration of IL22Fc 5 hours post *V. cholerae* infection was sufficient to sustain the body weight of infant mice and prolong their survival, suggesting that this treatment could be beneficial even after infection (Extended Data Fig. 6gh). Together, these findings suggest that IL22 may have clinical value in preventing or even treating cholera.

We also tested whether prophylactic IL22Fc could reduce the colonization of other enteric pathogens in infant mice. IL22Fc markedly limited *Citrobacter rodentium* colonization in the SI and colon (Extended Data Fig.6i). However, IL22Fc administration only led to a modest reduction in *Salmonella enterica* serovar Typhimurium intestinal colonization and did not reduce the pathogen burden in the liver or spleen (Extended Data Fig.6j), suggesting that the treatment did not influence pathogen dissemination. These results suggest that IL22Fc treatment is potent at impairing the intestinal colonization of extracellular pathogens in infant animals.

### Elevated requirement for *V. cholerae* motility in IL22 treated animals

To understand how IL22Fc-treatment alters the SI milieu, we compared *V. cholerae’s* genetic requirements for intestinal colonization in IL22Fc-treated and control mice using a transposon insertion site sequencing (Tn-seq) screen (Fig. 4a). In this genome-scale bacterial loss of function screen, comparisons of the abundances of transposon insertions in genes in the inoculum (input) to plated intestinal homogenates (output) are used to infer the genetic requirements for growth *in vivo*; reductions in the abundance of insertions in a gene in the output sample relative to the input suggests the gene promotes growth in the intestine. As expected, transposon insertions in genes encoding essential colonization factors, such as genes required for the biogenesis of the toxin-coregulated pilus (Tcp), were similarly depleted (compared to the inoculum) in both IL22Fc-treated and vehicle (PBS)-treated animals (Fig. 4b and Supplementary Table 6). In contrast, insertions in genes depleted more in IL22Fc-treated vs control animals compared to the inoculum represent genes that are more important for intestinal colonization in the IL22Fc-treated host. Notably, the relative depletion of insertions in 22 genes related to motility were 4-32x greater in IL22Fc-treated vs control animals (Fig 4b,c and Extended Data Table. 1), suggesting that motility is more important for *V. cholerae* colonization in IL22Fc-treated animals. Non-motile strains with deletions in genes important for flagellar assembly (*fliI* and *flgD*) and flagellar-based motility (*motB*)^29^ were created, and competition assays against wild-type strains corroborated the heightened importance of motility in IL22Fc-treated mice (Fig. 4d). We hypothesized that the increased importance of motility for colonization in the IL22Fc-treated mice reflects the role of motility in *in vivo* pathogen localization. Consistent with this idea, the fraction of luminal vs tissue-associated *V. cholerae* (Fig. 4e,f) was ∼86x greater in the IL22Fc-treated mice, demonstrating that IL22Fc treatment alters the intestine in a manner that impedes the pathogen from reaching the epithelial surface. In fact, it was difficult to detect fluorescently tagged *V. cholerae* closely associated with the epithelium in the crypts of treated mice (Fig. 4g). Together, these observations suggest that at least in part, IL22Fc treatment restricts *V. cholerae* colonization by limiting the pathogen’s access to the epithelial surface.

**Figure 4.**
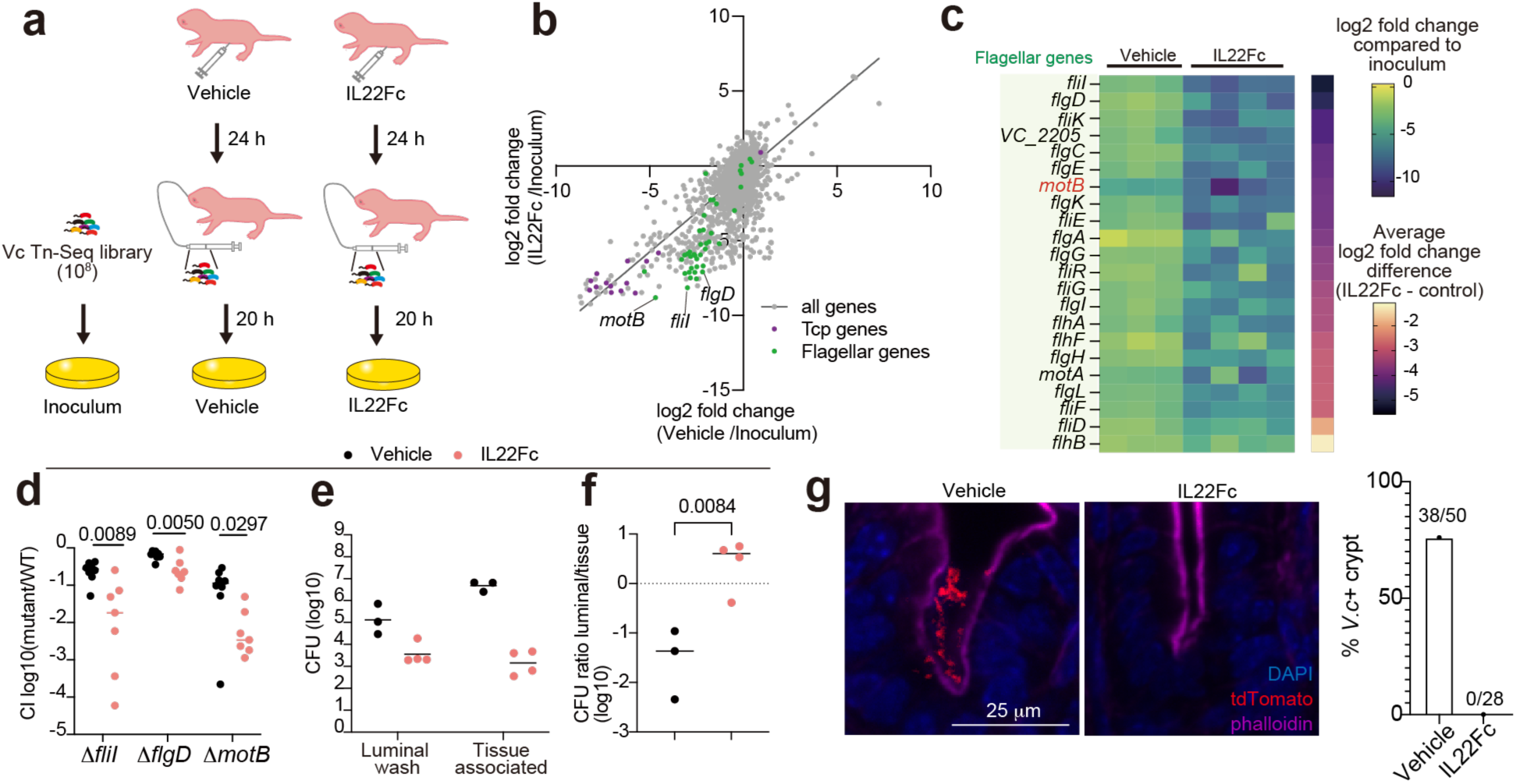
IL22Fc treatment modifies *V. cholerae* colonization requirements. **(a)** Experimental scheme of the transposon-insertion sequencing screen. Mice were given vehicle (PBS, n = 3) or 5μg of IL22Fc (n = 4) at 24 hours before infection and were infected with 10^8^ cells of a *V. cholerae* transposon library. SI tissues were harvested at 20 hpi. **(b)** XY plot of transposon insertion frequency compared to inoculum (culture on LB plates); x-axis represents the mean of log_2_ fold change (bacteria recovered from distal SI of PBS-treated mice vs. inoculum) and y-axis represents the mean of log_2_ fold change (bacteria recovered from distal SI of IL22Fc-treated mice vs. inoculum). Purple dots: Tcp-biogenesis mutants; green dots: Flagellar-related mutants. The plot represents the mean value of biological replicates (Vehicle: n = 3, IL22Fc: n = 4). (**c)** Heatmap representing log_2_ fold change of transposon insertion frequency in flagellar genes in samples recovered from mouse tissue compared to inoculum. Column and row show individual mouse and *V. cholerae* genes, respectively. The magma color heatmap indicates the difference in the average of log_2_ fold change between IL22Fc-treated mice vs. control mice. **(d)** Competitive index (CI) using barcoded mutant vs. WT strains in vehicle (black, n = 8) and IL22Fc-treated animals (red, n = 7). Multiple two-sided unpaired t-tests were performed between control vs IL22Fc-treated animals with adjustment using fold discovery rate. **(e**) CFU of *V. cholerae* detected in the luminal wash and attached to the tissue of the distal SI in vehicle-treated animals (black, n = 3) and IL22Fc-treated animals (red, n = 4). (**f**) Ratio of bacterial CFU in luminal washes compared with CFU of tissue-associated bacteria in vehicle-treated (black, n = 3) or IL22Fc-treated animals (red, n = 4). Two-sided unpaired t-test. Bars indicate the geometric mean and P-values are shown in individual figures (d-f). (**g)** Representative images of WT *V. cholerae* colonization in the crypt in control animal (left image) and IL22Fc-treated animal (right image), and a bar graph showing the percent of crypts occupied by *V. cholerae* in the distal SI. Blue: DAPI, red: *V. cholerae LacZ::tdTomato*, magenta: phalloidin for actin staining. Scale bar is shown.

### IL22 administration modifies the gene expression profiles and cellular composition of the SI epithelium

Treatment with IL22Fc significantly increased the length of the SI, but not the colon, without impacting the pups’ body weight (Extended Data Fig. 7a-c), suggesting that this cytokine modifies the development of the SI. 36 hours after IL22Fc administration, scRNA-seq of the SI epithelium revealed a marked increase in the abundance of cluster 3 and 9 cells (Fig 5a,b, Supplementary Table 1). Moreover, IL22Fc treatment stimulated changes in the transcriptomes of all types of epithelial cells in the distal SI (Extended Data Fig. 7d).

**Figure 5.**
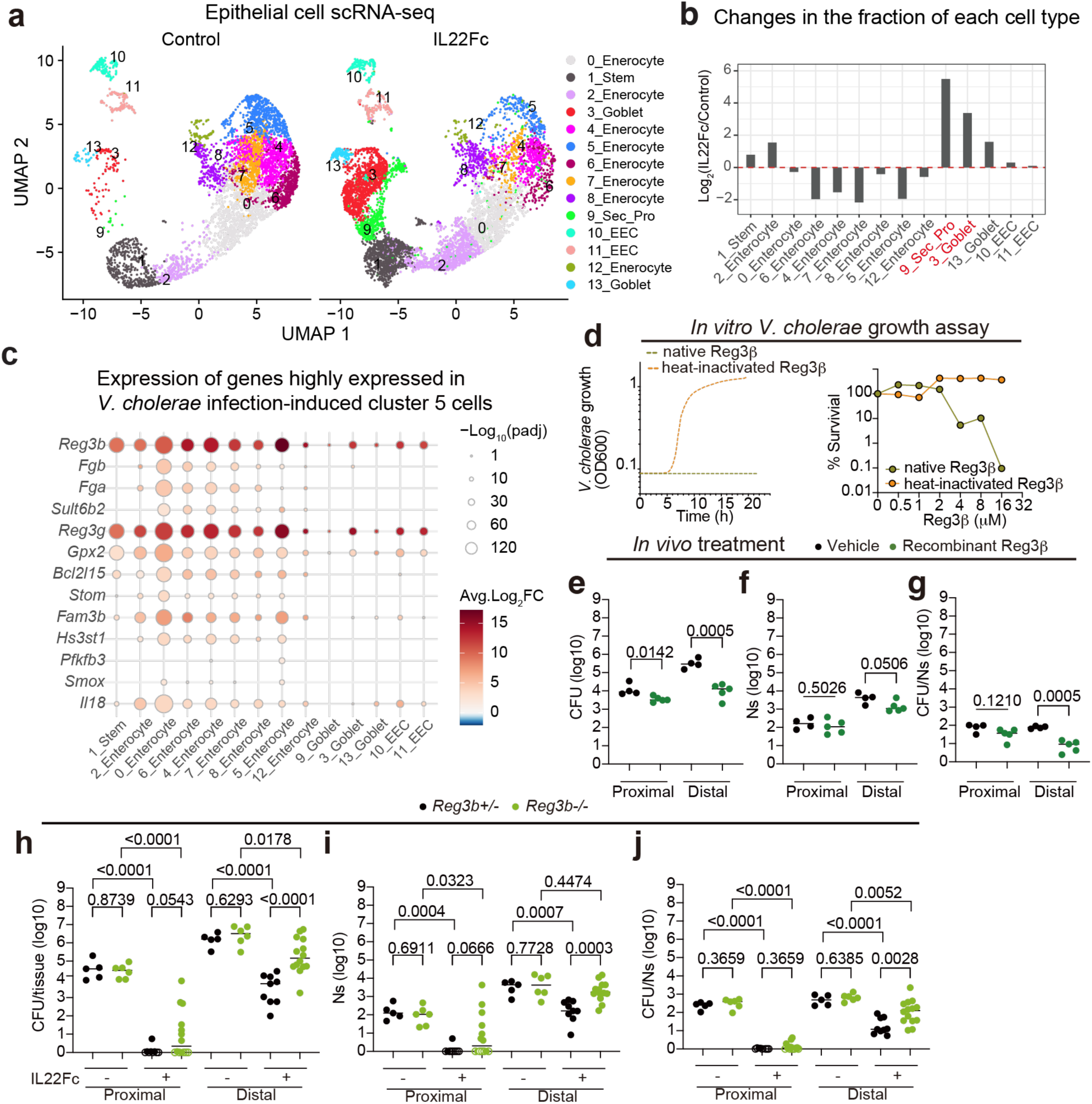
IL22Fc treatment alters intestinal epithelial cell transcriptional profiles. **(a)** UMAP of epithelial single-cell RNA-seq data of the distal small intestine in P5 infant mice in control and IL22Fc-treated animals. (**b)** Comparison of cluster abundance between IL22Fc-treated and untreated control animals. (**c)** Dot plot representing control vs. IL22Fc-treated gene expression changes of cluster 5 cell marker genes (Extended Data Fig. 3a) in epithelial cell subsets. Color represents average log_2_ fold change of gene expression compared to untreated control. Dot size represents negative log_10_-scaled adjusted p-value. Wilcoxon rank-sum test with Bonferroni correction. (**d)** *V. cholerae* growth curve measured by OD600 after incubation with native or heat-inactivated Reg3β (left). Percent *V. cholerae* survival after incubation with indicated concentration of Reg3β (right). Data represent mean for experimental replicates; native Reg3β (n = 4/condition), heat-inactivated Reg3β (n = 2/condition). (**e-g)** CFU (e), founding population (Ns) (f), and CFU/Ns (g) of *V. cholerae* in the proximal and distal SI at 16 hpi. Infant mice were orally administered 2 doses of vehicle (PBS, n = 4) or Reg3β recombinant protein (33 μg/dose, n = 5) at −0.5 and +1 hpi. Bars represent the geometric mean for biological replicates. One-way ANOVA. (**h-j)** CFU (h), Ns (i), and CFU/Ns (j) of *V. cholerae* in the proximal and distal SI of P5 *Reg3b*-/- mice and heterozygous littermate at 16 hpi. Bars represent the geometric mean for biological replicates; IL22Fc (-) *Reg3b^+/-^*: n = 5; IL22Fc (-) *Reg3b^-/-^*: n = 6; IL22Fc (+) *Reg3b^+/-^*: n = 9; IL22Fc (+) *Reg3b^-/-^*: n = 14. On circles indicate no CFU detected. One-way ANOVA. Sample groups were compared among the same tissue. P-values are shown.

Unexpectedly, IL22Fc treatment reduced the expression of more genes than it increased in 13 of 14 clusters (Extended Data Fig. 7d, Supplementary Table 7). There was a trend toward reduced expression of genes linked to lipid and sugar metabolism pathways as well as cell cycle genes in enterocytes of IL22-treated mice (Extended Data Fig. 7e-g), consistent with previous studies that elucidated a role of IL22 signaling in metabolism^30^. For some genes, IL22Fc administration induced changes in all cell types (e.g. increased expression of antimicrobial peptides) and in other cell subsets gene expression changes were more specific. For example, this treatment only stimulated expression of pathways associated with cell cycle (Mitosis, E2F, and MYC Target) in stem cells (Extended Data Fig. 7e). We confirmed stem cell proliferation in IL22Fc-treated animals with EdU staining (Extended Data Fig. 7h-j), as reported^31^. Notably, in most enterocyte clusters, genes highly expressed in cluster 5 enterocytes from *V. cholerae* infected animals were prominently upregulated by IL22Fc-treatment (Fig. 5c), suggesting that IL22 reprograms enterocytes to promote epithelial defense.

### IL22 induces expression of vibriocidal *Reg3b*

Among upregulated genes, pathway analysis revealed that IL22Fc increased expression of genes linked to synthesis of antimicrobial peptides in all epithelial populations (Extended Data Fig. 8a). *Reg3b* and *Reg3g* were among the most highly induced genes by Il22Fc treatment (Extended Data Fig. 8a); IL22Fc-treatment markedly elevated expression of *Reg3b* and *Reg3g* 10,000-fold compared to untreated animals, and in treated mice there was no further increase in expression of these antimicrobial proteins by infection (Extended Data Fig. 8b). The elevated expression of these antimicrobial proteins, which was also associated with IL22Fc treatment in a study of the colonic pathogen *Citrobacter rodentium* in adult mice^21^, along with *Defa24*, likely explains why antimicrobial pathways were one of the enriched pathways associated with this treatment in all clusters (Extended Data Fig. 8ab). However, proteomic analysis of mucosal washes from control and IL22Fc-treated animals only identified abundant Reg3β in treated animals, whereas Reg3γ, Defa17 and Defa24 were not identified (Extended Data Fig. 8c, Supplementary Table 8). Immunohistochemical staining confirmed that Reg3β is expressed all along the crypt-villus axis in IL22Fc treated animals (Extended Data Fig. 8d). Reg3β has been reported to kill several species of both Gram-negative^32^ and Gram-positive bacteria^33^ *in vitro,* but susceptibility of *V. cholerae* to this lectin has not been reported. In laboratory culture, we found that recombinant Reg3β decreased *V. cholerae* survival in a dose-dependent fashion, whereas heat-inactivated Reg3β failed to kill *V. cholerae* (Fig. 5d). Furthermore, oral administration of recombinant Reg3β protein decreased the *V. cholerae* burden as well as the size of the founding population and pathogen proliferation particularly in the distal SI (Fig. 5e-g), further confirming the vibriocidal activity of Reg3β *in vivo*. P5 *Reg3b^-/-^*mice were as susceptible to *V. cholerae* intestinal colonization as heterozygous littermates (Fig. 5h). However, IL22Fc administration to *Reg3b^-/-^* animals did not reduce the *V. cholerae* burden, size of the founding population, or amount of net proliferation as potently as in *Reg3b^+/-^* littermates (Fig. 5h-j), suggesting that the increased expression of Reg3β in IL22Fc treated animals contributes to the protective effects of IL22 *in vivo*.

Since *Reg3b^-/-^* animals were still protected from *V. cholerae* infection by IL22Fc treatment (Fig. 5h), other protective mechanisms beyond Reg3β induction likely exist.

### IL22 treatment increases the abundance of secretory epithelial cells

IL22 signaling is thought to be important for Paneth cell maturation and maintenance^34,35^. In P5 animals, where Paneth cells are absent (Extended Data Fig. 1d), IL22Fc administration increased the proportion of cells in the secretory lineage (cluster 3, 9, 13), especially cluster 3 (Extended Data Fig. 9a). Cells in clusters 3 and 9 express *Atoh1* and *Spdef*, key transcription factors for secretory lineage differentiation into goblet, Paneth, enteroendocrine, or tuft cells^36^. Based on marker genes, including *Muc2* and absence of *Lyz1* (Fig. 6ab), cluster 3 resembles adult goblet rather than Paneth cells^13,37^. However, cluster 9 has lower expression of goblet cell markers and expressed stem cell markers *Lgr5* and *Olfm4* (Fig. 6ab), suggesting it represents secretory progenitors^38^. Cells expressing *Atoh1* and *Spdef* reside primarily in crypts (Fig. 6c-e), and histologic assessment confirmed increased abundance of Muc2+ mucus-containing (PAS+) cells in the crypts of IL22Fc treated mice (Fig. 6f-h). Atoh1^Cre(ER-T2)^;Rosa26R^LSL-tdTomato^ mice, which encode Atoh1-dependent tamoxifen-inducible tdTomato, confirmed that crypt-associated mucous cells express *Atoh1* or derive from Atoh1+ cells (Extended Data Fig. 9b-d). To ask whether IL22Fc induces proliferation of secretory progenitor and goblet cells, we administered EdU and IL22Fc at the same time (Extended Data Fig. 9e). Compared with control animals, the numbers of EdU+ *Atoh1*+ cells were increased (Extended Data Fig. 9fg), indicating that IL22Fc triggers proliferation of *Atoh1*+ secretory progenitors or goblet cells. Together, these findings reveal that IL22Fc promotes secretory lineage and goblet cell development *in vivo*.

**Figure 6.**
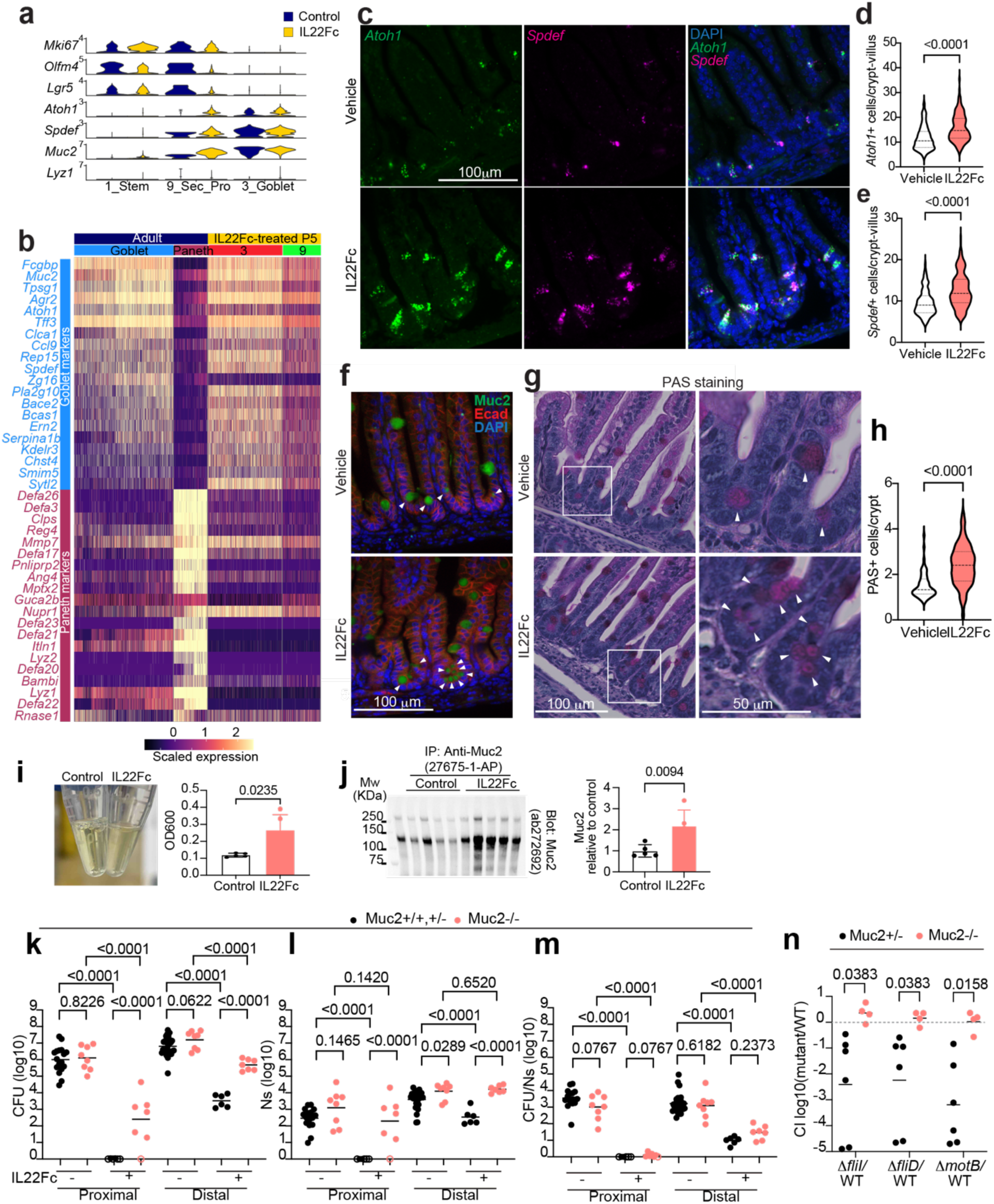
IL22Fc expands Goblet cells and Muc2 production is critical for IL22-mediated protection against *V. cholerae* colonization. **(a)** Violin plot of traditional marker gene expression in stem cells (*Lgr5* and *Olfm4*), proliferating cells (*Mki67*), secretory progenitor cells (*Atoh1* and *Spdef*), and goblet cells (*Muc2*) Paneth cells (*Lyz1*). (**b)** Heatmap of goblet and Paneth cell marker expression in adult (Haber et al: GSE92332^68^) and IL22Fc-treated P5 infant mice. (**c-e)** Representative RNA FISH images of *Atoh1* (green) and *Spdef* (magenta) expression and the number of *Atoh1*+ (d), or *Spdef*+ (e) cells in the epithelial cells in the crypts of the distal small intestine of vehicle- and IL22Fc-treated infant mice. Vehicle: n = 122 images from 4 animals; IL22Fc: n = 141 images from 4 animals. Two-sided unpaired t-test. (**f)** Representative images of immunohistochemistry of E-Cadherin (red), Muc2 (green), and DAPI (blue) in the distal small intestine of vehicle- and IL22Fc-treated animals. White arrowheads point to the Muc2+ cells in the crypts. Shown are representative of 41 images from 3 animals (vehicle) and 62 images from 3 animals (IL22Fc). (**g, h)** Representative PAS staining image (g) and enumeration of PAS+ cells (h) of the distal small intestine of vehicle and IL22Fc treated animals. Arrowheads indicate PAS+ cells in the crypt. 3 animals (Vehicle: n = 63 images) and 3 animals (IL22Fc: n = 65 images) were analyzed. Two-sided unpaired t-test with Welch’s correction. **(i)** Appearance of luminal washes collected from control and IL22Fc-treated infant mice (left). Turbidity of luminal washes were measured as OD600 in a spectrophotometer (right). n = 5/group. Two-sided unpaired t-test. Data are presented as mean values ± SD. **(j)** Immunoprecipitation-western blot for Muc2 in luminal washes (right blot) and the relative signal intensity to control group analyzed by densitometry (right bar graph). n =5/group. Two-sided unpaired t-test. Data are presented as mean values ± SD. (**k-m)** CFU (k), FP (l), and replication (CFU/FP) (m) of *V. cholerae* in the small intestine of P5 infant *Muc2*^-/-^ vs littermate wild type or heterozygous controls (*Muc2*^+/+, +/-^) animals treated with (+) or without (-) IL22Fc. Bars represent the geometric mean for biological replicates: *Muc2*^+/+, +/-^ IL22Fc (-) n = 28; *Muc2*^-/-^ IL22Fc (-) n = 8; *Muc2*^+/+, +/-^ IL22Fc (+) n = 6; *Muc2^-^*^/-^ IL22Fc (+) n = 7. One-way ANOVA. Sample groups were compared among the same tissue. P-values are indicated in individual plots. Open circles indicate no CFU detected. (**n**) Competitive index (CI) using barcoded mutant vs. WT *V. cholerae* strain in infant *Muc2*^-/-^ (n = 4, pink) or heterozygous littermate control (n = 6, *Muc2*^+/-^, black) mice treated with a single dose of 5 μg of IL22Fc 24 hours before infection. Bars represent the geometric mean for biological replicates. Two-sided multiple t-test with fold discovery rate. Corrected q-values are indicated in individual comparison.

Muc2 is the major gel-forming mucus protein expressed in goblet cells and secretory lineage cells in the SI^39^. IL22Fc-treatment increased the abundance of mucus and Muc2 protein in luminal contents (Fig. 6ij). Expression of glycosyltransferases, including *Mgat4c*, and *Fut2* were also induced by IL22Fc-treatment, as well by *V. cholerae* infection (Extended Data Fig. 10ab). Furthermore, modification of secreted Muc2 with fucose and *N*-acetylglucosamine was elevated in treated mice (Extended Data Fig. 10c). Thus, IL22Fc administration elevates the abundance of the Muc2+ cell population, as well as Muc2 secretion and modification.

To investigate if Muc2 plays a role in IL22 treatment mediated resistance to *V. cholerae* colonization, we compared the effects of IL22Fc administration on *V. cholerae* colonization in *Muc2^-/-^* and control (*Muc2^+/+^*or *Muc2^+/^*^-^) littermates. Without IL22Fc administration, *V. cholerae* colonization in *Muc2^-/-^* mice was comparable to that in control, with a modestly greater burden (2.4-fold) in the distal SI of *Muc2^-/-^* mice.

However, administration of IL22Fc had a much more potent effect limiting *V. cholerae* colonization in both the proximal and distal SI in control vs *Muc2^-/-^* mice; the *Muc2^-/-^* animals had a 817-fold (geometric mean) greater *V. cholerae* burden in the proximal SI and 146-fold greater *V. cholerae* burden in the distal SI than control littermates (Fig. 6k), suggesting that Muc2 contributes to the therapeutic activity of IL22Fc. Interestingly, the role of Muc2 in mediating the activity of IL22Fc appears to be primarily in limiting the pathogen’s capacity to establish a colonization niche in the intestine (i.e., its founding population size) rather than its proliferation (CFU/Ns) (Fig. 6lm). Since the Tn-seq screen (Fig.4a-c) suggested that *V. cholerae* motility becomes more important for colonization in IL22Fc-treated animals, we tested whether Muc2 impedes motility-dependent colonization. For these experiments, competition assays in IL22Fc-treated *Muc2^-/-^* mice were used to compare colonization of WT and motility-deficient strains *(τ1fliI*, *τ1fliD,* and *τ1motB*). In marked contrast to observations in *Muc2^+/-^*littermate controls, where the motility-deficient strains had a colonization defect, the motility-deficient strains colonized *Muc2^-/-^*as well as WT *V. cholerae* (Fig. 6n). Thus, the increased importance of flagellar-based motility for *V. cholerae* colonization in IL22Fc-treated animals (Fig. 4) is likely attributable to this cytokine’s stimulation of Muc2 production, which impedes pathogen access to its preferred niche along the epithelium in the crypts.

## Discussion

Given the extreme rapidity of cholera progression, innate defenses are particularly critical for protecting humans against *V. cholerae*. Here, scRNA-seq profiling of epithelial and lamina propria immune cell responses to experimental cholera in infant mice uncovered new facets of mucosal defense in the small intestine and suggested that IL22-dependent mechanisms play a role in cholera defense. IL22 therapy was highly protective against *V. cholerae* colonization and diarrhea in infant mice. scRNA-seq profiling of IL22-treated animals offered new insights into the mechanisms of this cytokine and uncovered an IL22-stimulated shift in the pattern of intestinal epithelial cell differentiation toward the secretory cell lineage. The products of these cells, including Muc2, appear to be critical for IL22-mediated protection from experimental cholera, a mechanism that may generally contribute to the therapeutic effects of IL22 (Extended Data Fig. 10d).

*V. cholerae* triggered changes in gene expression in most epithelial cell subsets but unexpectedly increased the abundance of an enterocyte subset (cluster 5) whose signature gene program was largely linked to defense. Since P5 mice lack Paneth cells, which are rich in antimicrobial peptides and are thought to protect the crypt stem cells^13^, cluster 5 enterocytes may function in an analogous fashion during early life before Paneth cells arise around P15^13^. However, in contrast to Paneth cells, whose relative abundance undergoes a modest increase during acute infection^40^, the abundance of cluster 5 enterocytes cells increased >10-fold during the 20 hours of exposure to *V. cholerae*. Because *V. cholerae* primarily localizes near the base of villi^14^, our findings raise the possibility that a consequence of the pathogen’s presence in the crypts is an alteration of the transcriptional program of adjacent enterocytes into this mucosal defense-associated epithelial subset. *V. cholerae* preferentially colonizes in the distal SI^14,41^ (Fig. 3bc). This localization may also explain why the epithelial and immune response is generally more pronounced in this part of the SI.

Several studies have revealed that IL22 deficient mice have heightened susceptibility to death from mucosal pathogens^20–22^. This work has led to the idea that IL22 promotes the integrity of the epithelium through a variety of mechanisms, including increases in antimicrobial proteins^21^, epithelial cell surface modifications^42^, and Paneth cell differentiation^34^. For enteric pathogens, such as *C. rodentium* and *C. difficile*, experimental infections in *IL22*-/- mice have not revealed marked increases in intestinal pathogen burdens^21,22,43^, and heightened mortality has been attributed to dissemination of intestinal bacteria, at least in part due to reduced activity of complement C3^22^. Fewer studies have examined the consequences and the mechanisms of IL22 administration on pathogen burden. In adult mice *C. rodentium* infection, prophylactic administration of IL22Fc only modestly reduces colonic colonization, but significantly reduces disease by preventing epithelial damage^44^.

In contrast, administration of IL22Fc almost entirely ablated the susceptibility of P5 mice to *V. cholerae* colonization. Several IL22-dependent mechanisms likely account for the marked reduction of *V. cholerae* colonization in IL22Fc-treated animals. Although there was a modest benefit to administering IL22Fc after infection, the most dramatic therapeutic effects of IL22Fc required that the cytokine be administered 24-48 hours before pathogen inoculation, indicating that time is required for the beneficial effects of IL22Fc to accrue. IL22Fc stimulated alterations in the transcriptional profile, cellular composition and even the length of the SI in P5 animals. One of the striking changes elicited by IL22Fc administration was the augmentation in expression of antimicrobial genes in most enterocyte subsets. For example, Reg3β became detectable in epithelial cells throughout the crypt-villus axis. However, since we found that the absence of Reg3β, in *Reg3b^-/-^* animals, did not render mice more susceptible to *V. cholerae* colonization, physiologic levels of Reg proteins may be insufficient to impede colonization. Instead, the supraphysiologic levels of Reg3β proteins and potentially other antimicrobial products induced by IL22Fc treatment likely are important for the beneficial effects of IL22. Similarly, IL22Fc administration markedly diminished colonic colonization with *C. rodentium* in infant mice (Extended Data Fig. 6i), even though *Reg3b^-/-^* and *Reg3g^-/-^* mice do not exhibit elevated pathogen burden or disease severity associated with this pathogen in an adult mouse model^43,45^.

Although elevated Reg3β contributes to IL22Fc-mediated protection, our data suggests that the major mechanism underlying reduced *V. cholerae* colonization in IL22Fc-treated animals is due to elevated Muc2 production. IL22Fc administration appears to drive the differentiation of stem cells towards production of secretory cells *in vivo*, as suggested in organoids^35^. Studies in suckling *Muc2^-/-^* mice revealed that IL22Fc administration was not nearly as potent at impeding *V. cholerae* intestinal colonization as in littermate *Muc2^+/-^*controls. Thus, mucin production appears to be required for robust IL22Fc activity. We propose that the increase in goblet cell number and Muc2 secretion in IL22Fc treated animals blocks the pathogen from reaching its preferred niche, where it establishes a founding population in close apposition to the epithelium of the crypt^41,46^. The IL22Fc-stimulated impedance of pathogen access to the epithelium and crypts were apparent in the results of our TnSeq screen (Fig. 4). Moreover, the competitive advantage of motility for colonization in IL22Fc-treated animals was not observed in *Muc2*^-/-^ mice (Fig. 6n), demonstrating the linkage between mucus-mediated protection and *V. cholerae* motility. Glycosylation of Muc2 may also modulate disease outcome by reducing cholera toxin expression^47^. Finally, since mucus is decorated with glycans and antimicrobial proteins, and their functions are entwined^48^, the dramatic beneficial effects of IL22Fc on experimental cholera likely derive from synergistic interactions between these components of innate intestinal defense. Overall, the marked reduction in *V. cholerae* colonization likely explain the IL22Fc-mediated protection from cholera-like death in suckling mice.

Given the dramatic benefits of IL22Fc in the suckling mouse model of cholera and the emergence of highly antibiotic-resistant *V. cholerae*^49^, studies of this fusion protein as a prophylactic agent for individuals susceptible to household contact infection^50^ and/or as an adjunctive therapy along with fluid resuscitation for cholera should be considered. Moreover, IL22Fc administration may be an important therapeutic adjuvant for other extracellular enteric pathogens such as enterotoxigenic *E. coli,* another pathogen that like *V. cholerae*, does not damage the epithelium. However, several issues could complicate the translation of our findings to humans. First, although infant mice represent an outstanding model for studies of *V. cholerae* intestinal colonization^12^ , these animals differ from humans in several important regards. For example, human Paneth cells develop during embryogenesis, whereas mouse Paneth cells develop around postnatal day 15^13,37^. Furthermore, the mechanisms by which IL22 protects the host from infection may differ according to age, since IL22Fc administration did not elevate goblet cell numbers in adult mice^34^ unlike our findings in infant mice. Variations in epithelial maturation, including the expression levels of IL22RA1/IL10RB, the IL22 receptor, in small intestinal epithelial cells could impact on the efficacy of IL22Fc.

Despite these limitations, several clinical trials have already revealed the safety of IL22Fc in humans. The primary dose-limiting adverse events when IL22Fc was administered repeatedly were dermatologic^27,51^. Many additional studies, including whether IL22Fc administration would be beneficial in the epidemiologic conditions, such as undernutrition where cholera is prevalent, will be necessary to establish the safety of IL22Fc in relevant populations. Costs of this fusion protein could be prohibitive but alternative cheaper methods to promote endogenous IL22 production such as tryptophan metabolites that promote ILC3 differentiation and IL22 production^52^ may be feasible.

## Materials and Methods

### Mice

C57BL/6NCrl (strain code: 027) were purchased from Charles River Laboratory. ROSA26LSL-TdTomato (B6.Cg-Gt(ROSA)26Sortm9(CAG-tdTomato)Hze/J; RRID:IMSR_JAX:007909), Lgr5-EGFP-IRES-creERT2 (B6.129P2-Lgr5tm1(cre/ERT2)Cle/J; RRID:IMSR_JAX:008875), IL22 knockout (C57BL/6-Il22tm1.1(icre)Stck/J; RRID:IMSR_JAX:027524) mice were from Jackson Laboratory (Bar Harbor, ME), *Atoh1^Cre(ER-T2)^* mice were a kind gift from Mikio Hoshino^53^, and *Muc2* knockout mice are gift from Anna Velcich^54^. *Reg3b* knockout (C57BL/6NCrl-Reg3bem1(IMPC)Mbp/Mmucd, RRID:MMRRC_067360-UCD) was obtained from the Mutant Mouse Resource and Research Center (MMRRC) at University of California at Davis, an NIH-funded strain repository, and was donated to the MMRRC by Kent Lloyd, D.V.M., University of California, Davis. Mice were bred in the Bio Safety Level 1 (BSL1) animal facility in Brigham and Women’s Hospital.

Mice were maintained in a BSL1 animal facility at Harvard Medical School or Brigham and Women’s Hospital and transferred to the BSL2 animal facility for the infection experiments. Mice were kept on a 12-h light/dark cycle at 20–24 °C with 50% humidity. Animals were maintained in cages filled with pine wood bedding and paper nesting material with chow (LabDiet 5053) and water by the animal facility staff. All animal experiments were performed according to protocols reviewed and approved by the Brigham and Women’s Hospital Institutional Animal Care and Use Committee (protocol 2016N000416) and in compliance with the Guide for the Care and Use of Laboratory Animals. Both male and female mice were mixed and randomly assigned to experimental groups.

### Bacteria Strains and culture

A *Vibrio cholerae* O1 Ogawa clinical isolate from the 2010 Haiti outbreak (HaitiWT)^55^ was used here. Motility deficient mutants *(τ1motB, τ1flgD, τ1fliI*) were created using homologous recombination. To create targeting vectors, 1000 bp of DNA flanking each side of the targeted genes were amplified with Phusion DNA polymerase (NEB) and these segments were cloned into pCVD442’s SmaI site using NEBuilder HiFi DNA assembly kit (NEB).

Barcoded WT or motility deficient mutants *V. cholerae* were generated as previously described^56^. Fluorescent protein tdTomato labeled HaitiWT was created previously^57^. Primers used for the plasmid construction are listed in supplementary data (Supplementary Table 9). A streptomycin resistant isolate of *Citrobacter rodentium* strain ICC168 and *Salmonella Enterica* Serovar Typhimurium SL1344 strain were also used.

### *In vivo* infection and colony forming unit (CFU) counting

*V. cholerae* was cultured in lysogeny broth (LB) with streptomycin (Strep) 200 μg/mL at 37 °C overnight (∼18 h) prior to animal experiments. Mice at postnatal day 5 ± 1 (P5) (2.5-3.5 g of body weight, sex mixed) were infected with 1×10^5^ Vc^STAMP^ by oral gavage using PE-10 tubing. Weight and survival were monitored every 4-8 hours, and animals were euthanized when they lost 20% of body weight. 3 cm of proximal SI (samples were dissected after the gastro-duodenal junction) and distal SI (∼3 cm from the end of the ileum) were homogenized with stainless beads in 33% glycerol LB. The homogenates were serially diluted and plated on LB Strep agar plates for CFU counting.

*C. rodentium* was cultured in LB media containing 200 μg/mL strep at 37 °C overnight (∼24 h). P5 infant mice were infected with 1×10^8^ cells of *C. rodentium* by oral gavage using PE-10 tubing. Whole SI, and colon were dissected and homogenized in PBS and serial dilutions of the tissue homogenates were plated on LB Strep agar plates for CFU counting.

*S.* Typhimurium was cultured in LB Strep 200 μg/mL at 37 °C overnight prior to animal experiments. P5 infant mice were infected with 1×10^7^ cells of *S.* Typhimurium by oral gavage using PE-10 tubing. Infant mice were euthanized 1- or 3-days post infection. Whole SI, colon, liver, and spleen were dissected and homogenized in PBS, and serial dilutions of the tissue homogenates were plated on LB Strep agar plates for CFU counting.

### CD45+ immune cell isolation from small intestine lamina propria

Infant mice were euthanized with isoflurane. The proximal and distal SI, as described above, were collected, cut open longitudinally, and stored separately in buffer [2% Fetal Bovine Serum (FBS), 50 μg/mL Gentamicin (Gen), Penicillin-Streptomycin (PS) PBS] on ice until all animals were dissected. Tissues were incubated in 10 mL of reducing buffer [10 mM DTT 10 mM HEPES 2% FBS, 50 μg/mL Gen, PS, RPMI 1640], for 10 min on ice, followed by incubation in 10 mL of epithelial dissociation buffer [5 mM EDTA, 10 mM HEPES, 2% FBS RPMI1640] at 37℃ for 25 min. The remained tissue was washed in PBS, cut into small pieces, and incubated in digestion buffer [0.5 mg/mL collagenase D, 0.3 mg/mL Dispase II, 0.05 mg/mL DNase I, 10 mM HEPES 5% FBS, RPMI 1640] at 37℃ for 30 min. After vortexing for 20 sec, dissociated tissue was passed through a 40 μm filter and centrifuged at 300 xg for 5 min. The immune cells were further separated in a 40% /80% Percoll gradient by centrifugation at 1900 xg for 30 min. The layer between 40% and 80% Percoll was collected, mixed with FACS buffer [2% FBS 2mM EDTA PBS], and centrifuged at 300 xg for 5 min. Cells were resuspended in FACS buffer containing anti-CD16/32 antibody, and stained with anti-CD45 FITC antibody. After being washed in FACS buffer, cells were stained with propidium iodide before sorting, and the CD45+ PI- population was isolated using a cell sorter (SH800Z, Sony, Japan).

### Single-cell isolation of Epcam+ epithelial cell

Infant mice were orally infected with 10^5^ WT *V. cholerae* cells and euthanized 20 hpi. For IL22Fc (Genentech)^27^ treatment, P4 mice were administered 20 μg of IL22Fc intraperitoneally and returned to their dam. After 36 hours, the mice were euthanized. Proximal and distal SI were collected, cut open longitudinally, and washed in Gut wash media [2% FBS, 10 mM HEPES, 100 μg/mL Pen/Strep, 50 μg/mL gentamicin in RPMI1640] in 15 mL conical tubes. Tissues were rinsed in ice-cold PBS, incubated in Epithelial dissociation solution [30 mM EDTA in PBS] on-ice for 40 min, shaken gently 30 times every 5 min, and then vortexed for 20 seconds. Tissue pieces were removed with forceps. Dissociated cells in the supernatant were centrifuged and resuspended in ice-cold PBS. This step was repeated twice to wash out EDTA. The cell pellets were resuspended in Dispase solution [2% FBS, 0.1 mg/mL dispase in 4 mL RPMI1640 per animal] at RT for 3-5 min, then cells were dissociated by pipetting with a wide-bore tip, followed by pipetting 10 times with normal tips. 4 mL of ice-cold RPMI1640 containing 2% FBS was added to the cell suspension and filtered through a 70 μm cell strainer. The strainer was washed with ice-cold RPMI1640 containing 2% FBS to collect the cells. Cells were pelleted by centrifugation, stained with anti-Epcam FITC (1:100), and passed through a 100 mm cell strainer. After being stained with LIVE/DEAD Fixable Violet Dead cell staining kit (ThermoFisher, L34964), live Epcam+ cells were sorted by a SONY SH800Z Cell sorter.

### Single-cell RNA-Seq

Sorted CD45+ SI lamina propria (LP) immune cells or Epcam+ epithelial cells were collected into PBS containing 2% FBS, followed by centrifugation at 1000 xg for 5 min, resuspended in PBS, counted, and passed through a 40 μm cell strainer. Single-cell suspensions were applied to a 10X Chromium Controller for GEM generation using Chromium Next GEM Single Cell 3’ Reagent Kits v3.1 (10x Genomics). After barcoded cDNA generation, samples were amplified with the index primers (Dual Index Kit TT Set A, 10x Genomics) for 12 cycles. The libraries were sequenced on a NovaSeq 6000 S1 or NextSeq500. Data analysis was performed using 10x Cloud Analysis, and the filtered feature H5 files were used for further analysis with the Seurat package in R. Seurat objects were generated using Seurat v5.0.0 (https://satijalab.org/seurat/). Low-quality droplets (nFeature_RNA > 200 & nCount_RNA > 1000 & percent.mt < 20) and potential multiplets detected by scDblFinder were removed. Data from uninfected and infected proximal and distal SI samples were normalized by SCTransform (method = “glmGamPoi”, vst.flavor = “v2”) and integrated by Seurat’s functions. Clustering was performed with Seurat’s FindCluster (dims = 1:30, resolution = 0.4 for epithelial, 1.0 for CD45+ cells).

Marker genes and differential gene expression analysis on each cell cluster was performed using the FindMarkers function of Seurat, and we annotated clusters based on canonical marker genes. Single-cell pathway analysis (SCPA) was performed by SCPA package v1.6.1 with default settings (downsample = 500, min_genes = 15, max_genes = 500) for epithelial cells, and downsample = 300 for CD45+ immune cells. Pathway enrichment was obtained by multiplying −1 to the fold change in the SCPA output. Pathways containing the following words were selected for coloring in Fig.1d: “INTERFERON” for IFN; “TNFA”, “INFLAMMATORY”, “INFLAMMATION” for TNFα, Inflammatory; “M_PHASE”, “S_PHASE”, “PRO_PHASE”, “ANAPHASE”, “CHROMATIDS”, “G2_M_CHECKPOINTS”, “SYNTHESIS_OF_DNA”, “MITOTIC”, “DNA_REPLICATION”, “MYC”, “E2F” for Mitosis; “TRANSLATION”, “RIBOSOME”, “RIBOSOMAL” for Translation and Ribosome; “GOLGI”, “MTORC”, “UNFOLDED”, “HYPOXIA” for Golgi, ER, mTOR; “AMINO_ACID”, “AMINO_ACIDS”, “FATTY_ACID”, “OXIDATIVE_PHOSPHORYLATION”, “TCA_CYCLE” for Metabolism. For Fig.2b: “INTERFERON” for IFN; “IL17”, “IL23”, “PROINFLAMMATORY” for Proinflammatory, IL17, IL23; “TRANSLATION”, “RIBOSOME”, “RIBOSOMAL” for Translation, Ribosome; “GOLGI”, “UNFOLDED”, “MTOR”, “HYPOXIA” for Golgi, ER, and mTOR. For Extended Data Fig.7e: “ANTIMICRO” for Antimicrobial peptides; “M_PHASE”, “S_PHASE”, “PRO_PHASE”, “ANAPHASE”, “CHROMATIDS”, “G2_M_CHECKPOINTS”, “SYNTHESIS_OF_DNA”, “MITOTIC”, “DNA_REPLICATION” for Mitosis; “TRANSLATION”, “RIBOSOME”, “RIBOSOMAL” for Translation and Ribosome; “E2F” for E2F; “MYC_” for MYC Target; “AMINO_ACID”, “AMINO_ACIDS”, “LIPID”, “GLUCOSE”, “SUGAR”, “FATTY” for Lipid, Sugar, Amino acid. HALLMARK and KEGG pathways containing the following words were plotted in Extended Data Fig. 2c: “AMINO_ACID”, “AMINO_ACIDS”, “FATTY_ACID”, “OXIDATIVE_PHOSPHORYLATION”, “TCA_CYCLE”.

In RNA velocity analysis, *V. cholerae*-infected intestinal epithelial cells scRNA-seq data were processed using Cell Ranger, velocyto, Seurat and scVelo, according to the scVelo instruction (https://scvelo.readthedocs.io/en/stable/index.html). Briefly, spliced and unspliced transcript counts were quantified from Cell Ranger output using velocyto^58^ (v0.17.17) with the mouse reference genome (mm10). To maintain consistency with our prior single-cell analysis, velocity data was integrated with the Seurat (v5)-processed dataset. Cell barcodes from velocyto loom files were matched to the quality-controlled Seurat object (after doublet removal and QC filtering). Pre-computed UMAP coordinates and cluster annotations from the Seurat SCTransform/RPCA integration workflow were imported into the scVelo analysis framework, ensuring identical cell embeddings between the two analyses. RNA velocity was estimated using scVelo^59^ (v0.3.0) in Python (v3.10) and velocity streamlines were projected onto the UMAP embedding to visualize cellular state transitions.

Cell-cell communication analysis between epithelial and immune cells were performed using the CellChat^19^ (v2.2.0). First, epithelial and immune cell single cell RNA-seq data were integrated according to condition (uninfected or infected). Cellchat objects were generated using createCellChat function. Comparison of the communication probabilities mediated by ligand-receptor pairs from LTi-like ILC3 to enterocytes were performed by netVisual_bubble function and visualized by netVisual_chord_gene function with default settings.

### Flow cytometry

SI lamina propria cells were isolated according to the protocol above. For intraepithelial lymphocyte (IEL) isolation, distal SI tissue were cut open longitudinally and incubated in 10 mL of reducing buffer [10 mM DTT 10 mM HEPES 2% FBS, 50 μg/mL Gen, PS, RPMI 1640], for 10 min on ice, followed by incubation in 10 mL of epithelial dissociation buffer [5 mM EDTA, 10 mM HEPES, 2% FBS RPMI1640] at 37℃ for 25 min. After removal of the remaining tissue pieces by pipetting, supernatants were passed through a 70 μm cell strainer and spun down at 300 xg for 5 min. The pellets were resuspended in 2 mL FACS buffer and overlayed onto 40%/80% Percoll gradients. Cells accumulated between 40% and 80% Percoll were transferred to new tubes, washed in 5 mL of FACS buffer, and centrifuged at 500 xg for 5 min. Cell pellets were washed in 2% RPMI1640 media and used for cell culture.

Cells isolated from LP and IEL were cultured in 10 % FBS RPMI1640 supplemented with 1x Brefeldin A (Biolegend) and Penicillin/Streptomycin (Gibco) for 3 h. Cells were mixed with anti-CD16/32 antibody (Biolegend, 101302) to block Fc binding, and stained with the following combination of antibodies in 2% FBS 2mM EDTA PBS; Live/Dead Aqua (ThermoFisher, L34957), anti-CD45+ Alexa Fluor 700 (BD Biosciences, 560510), anti-CD3e Super Bright 600 (ThermoFisher, 63-0031-82), anti-B220 FITC (Biolegend, 103206), anti-CD11b FITC (Biolegend, 101206), anti-CD11c FITC (BD Biosciences, 557400), anti-TCRγ/8 APC (Biolegend, 118143). After 30 min incubation at room temperature, cells were washed in 2% FBS 2mM EDTA PBS and fixed in Fixation/ Permeabilization solution (ThermoFisher, 88-8824-00) at 4 °C overnight. After washing, cells were stained with anti-IL22-PerCP-eFluor710 (ThermoFisher, 46-7222-80) at room temperature for 30 min and analyzed using a Cytek Northern Light (Cytek, MA).

### Immunohistochemistry and histology

Distal small intestines were fixed in 4% paraformaldehyde (PFA) PBS for 4 hours and dehydrated in 20% sucrose-PBS overnight. Tissue was embedded in OCT Compound: 20% sucrose-PBS = 2.5:1 and frozen by liquid nitrogen. Tissue blocks were cut into 10 μm sections with a cryostat at −18 °C (Leica) and the sections were dried at room temperature for 2h and stored at −80 °C until use. Tissue sections were washed in PBS, blocked in blocking buffer [5% goat serum 0.1% Triton X100 PBS] for 30 min, and stained with sheep anti-Reg3β antibody (1:100, R&D Systems), rabbit anti-*V. cholerae* antisera Ogawa (1:100, BD Difco) at 4 °C overnight. Tissue sections were washed in PBS twice and stained with anti-sheep IgG Alexa Flour 555 (1:200, ThermoFisher), goat anti-rabbit IgG Alexa Flour 488 (1:200, ThermoFisher), or chicken anti-rabbit IgG Alexa Flour 647 (1:200, ThermoFisher) and DAPI (1:1000, ThermoFisher) at room temperature for 30 min. The sections were washed in PBS and covered with Prolong gold antifade reagent (ThermoFisher).

For PAS staining for paraffin sections, fixed tissue was processed, sectioned, and stained at the Harvard Rodent Histology core. For Muc2 staining, paraffin sections were deparaffinized with xylene for 10 min x2, rehydrated in 100%, 80%, 70%, 60% ethanol, and PBS for 5 min in each solution. Sections were incubated with antigen retrieval buffer (citrate buffer, pH 6.0) and microwaved for 10 min. Rehydrated tissue was blocked in blocking buffer for 30 min, stained with rabbit anti-Muc2 antibody (1:100, Proteintech) in blocking buffer overnight, washed in PBS, and stained with anti-rabbit IgG Alexa Flour 488 (1:200, ThermoFisher) at room temperature for 30 min. The tissue was stained with DAPI, washed in PBS, and mounted in Prolong gold antifade reagent (ThermoFisher, P36934).

Fluorescent and PAS stain images were imaged with a Nikon Eclipse microscope with a Nikon Microscope Objective Plan Apo l 20x/0.75 OFN25 DIC N2 lens, Plan Flour 40x/1.30 Oil lens, or Plan Apo VC 100x/1.40 Oil DIC lens (Nikon) and captured with a Andor iXon DU-897E-CS0-#BV-500 Camera (Oxford Instruments).

### Single molecule RNA Fluorescence *in situ* hybridization (RNAScope)

For detection of *Fgb* and *Sult6b2*, the dissected distal SI tissue was fixed in 4% PFA-PBS overnight at 4°C and dehydrated in 20% Sucrose-PBS overnight. Tissue was embedded in OCT Compound: 20% sucrose-PBS = 2.5:1 and frozen by liquid nitrogen. Tissue blocks were cut into 10 μm sections with a cryostat at −20 °C (Leica). The sections were attached to MAS coated slide glass (Matsunami), dried at −20°C for 2h and were stored at −80 °C until use. Tissue sections were stained according to the manufacture’s RNAScope protocol (ACDbio) with hybridization probes: Mm-Sult6b2-C3 (1789851-C3) and Mm-Fgb-C2 (578921-C2). For detection of *Atoh1* and *Spdef*, P4 infant animals were administered PBS or 20 μg of IL22 and were returned to their dam. After 36 h, mice were euthanized, and tissues were processed as described above. Tissue sections were stained with hybridization probes Mm-Atoh1 (408791) and Mm-Spdef-C2 (544421-C2). Stained sections were imaged using a Nikon Eclipse microscope as above.

### Proliferating cell staining with 5-ethynyl 2’-deoxyuridine (EdU)

For the staining of *V. cholerae* infected animal tissues, P5 animals were randomly assigned to receive 40 μg of EdU at 0 hpi or 18 hpi. The mice were infected with 10^5^ cells of *V. cholerae* and euthanized and dissected at 20 hpi. For the staining of IL22Fc-treated animal tissues, P4 infant littermate mice were randomly assigned to receive either PBS or IL22Fc. Mice were administered 40 μg of EdU and PBS or 20 μg of IL22Fc intraperitoneally and returned to their dam. After 36 hours, the mice were euthanized and dissected. For both experiments, the SI tissue was fixed, embedded, and cut into 10 μm cryosections as described above in methods for Immunohistochemistry or RNAScope. After the immunohistochemistry or RNAScope staining procedure, tissue sections were stained with Click-iT™ EdU Cell Proliferation Kit for Imaging, Alexa Fluor™ 647 dye (ThermoFisher), according to the manufacture’s protocol. Figure images were captured with Nikon Eclipse microscope as above.

### Lineage tracing

*Lgr5^Cre(ER-T2)^;Rosa26R^LSL-tdTomato^*mice were given 200 μg of tamoxifen 9 hours before and again at the same time as infection with 10^5^ cells of *V. cholerae*. Mice were euthanized at 20 hpi. The SI was fixed, embedded, and cut into 10 μm cryosections as described above (See RNAScope). Tissue sections were stained with hybridization probes: Mm-Sult6b2-C3 (1789851-C3) and Mm-Fgb-C2 (578921-C2) for RNAScope and further stained with anti-RFP antibody (1:200, Rockland Immunochemicals, 600-401-379). After washing in PBS, tissue was stained with chicken anti-rabbit IgG Alexa 647 (1:200, ThermoFisher, A-21443) for 30 min, stained with DAPI (1:500) for 5 min, and mounted in Prolong Gold Antifade Reagent.

*Atoh1^Cre(ER-T2)^;Rosa26R^LSL-tdTomato^* mice were given 200 μg of tamoxifen 9 hours before and again at the same time as IL22Fc (20 μg), and euthanized 37 hours after IL22Fc administration. The tissue was processed according to the protocol for immunohistochemistry (See above). For Ki67 staining, rabbit anti-Ki67 antibody (1:100, Abcam, ab16667) and anti-rabbit IgG Alexa 647 (1:200) were used. For mucus positive secretory lineage staining, tissue section was stained with UEA-I FITC (1:100, vector), phalloidin Alexa 647 (1:200, ThermoFisher), and DAPI (1:1000), for 10 min, washed in PBS, and mounted in Prolong Gold Antifade Reagent. Figure images were captured with Nikon Eclipse microscope as above.

### iDISCO clearing and 3D imaging

Two-centimeter pieces of distal small intestine were harvested from uninfected and infected animals and fixed in 4% paraformaldehyde PBS at 4°C overnight. Tissues were washed in PBS with 0.2% Tween-20 and 100 μg/mL Heparin for 1 hour twice and dehydrated gradually in 20%, 50%, 80% methanol (in PBS) for 1h, and twice in 100% methanol for 1h. Samples were then incubated overnight at room temperature on a shaker (50-60 rpm) in 66% Dichloromethane (Sigma) /33% Methanol. Samples were then incubated with 5% H_2_O_2_ in 20% DMSO at 4’C overnight, and then washed in 100% methanol, then 20% DMSO/methanol, and then rehydrated with 80%, 60%, 50%, 40%, 20% methanol in PBS for 1h each, and PBS for 1h twice. Tissues were then permeabilized in 0.2% Triton X-100 at RT for 1h, and 0.2% Triton X-100/20% DMSO/0.3M Glycine for 1day at 37°C with gentle agitation. After being washed in PBS with 0.2% Tween-20 and Heparin (100 μg/ml) at RT for 1h twice, tissues were blocked in PBS with 0.2% TritonX-100/10% DMSO/6% Donkey Serum at 37°C with gentle agitation for 1 day, stained with rat anti-IL22 (1;100. Thermofisher) in PBS with 0.2% TritonX-100/10% DMSO/3% Donkey Serum at 4 °C for 2 days. The tissues were washed in wash buffer [PBS with 0.2%Tween-20, 100 μg/mL Heparin] for 30 min for 5 times, incubated with secondary antibody; anti-rabbit IgG Alexa 647 (1:200, Invitrogen), anti-rat IgG Alexa 568 (1:200, Invitrogen), and rabbit anti-Epcam Alexa 488 (1:100, Biolegend) in PBS with 0.2% TritonX-100/10% DMSO/6% Donkey Serum at 37C for 1 day. Samples were washed in wash buffer for 5 times, stained with 2μg/mL DAPI in PBS, washed in wash buffer for 5 times. To clear the tissue, samples were dehydrated with 20, 40, 50, 80, and 100% methanol and incubated in 66% DCM/33% methanol at room temperature for 3 hours, subsequently incubated in 100% dichloromethane at room temperature for 15 min twice, and incubated in 100% ethyl cinnamate. Tissues were placed on a glass bottom dish, and covered with100% ethyl cinnamate. Three dimensional images were captured using a confocal microscope (Stellaris 8 Falcon DIVA, Leica) with Fluotar 16x/0.6 multi-immersion lense, and LasX acquisition software, built with arivis Vision 4D software version 4.1.2 (Zeiss).

### Quantitative PCR

Distal SI tissue from animals uninfected or infected with WT *V. cholerae* was collected at 20 hpi and homogenized in Trizol (ThermoFisher) with a bead beater with 3.2 mm metal beads. Total RNA was isolated according to the manufactures’ protocol. Total RNA was used for One-step cDNA synthesis and qPCR with the Luna Universal One-Step RT-qPCR Kit (NEB, E3005L) according to the manufacture’s protocol. The amplification signal was analyzed using a StepOnePlus Real-Time PCR System (Applied Biosystems). qPCR primer pairs are listed in Supplementary Table 9.

### Sequence tag-based analysis of microbial population dynamics (STAMP)

Infant mice were orally infected with barcoded Vc^STAMP^ and 16-20 hours later sacrificed and tissue homogenates were plated for CFU counting as described above. The remaining SI homogenates were plated on large (15cm diameter) LB-streptomycin plates and after overnight culture at 37°C, cells were collected in 33% glycerol LB and stored at −20 °C. Two microliter of each cell suspension was diluted with 100 μL milliQ water and boiled at 95 °C for 15 min. Each sample was amplified with One Taq DNA polymerase, forward and reverse primers with unique sequences that enable samples to be distinguished were used. The amplicons were purified using the Genejet PCR purification kit and sequenced with NextSeq P1 100 cycle Reagent Kit (Illumina). The barcode sequences were analyzed with STAMPR pipeline as described^28^.

### Transposon insertion sequence (Tn-seq)

A Tn-seq library in *V. cholerae* strain HaitiWT was used^60^. The library contained ∼195,000 distinct transposon insertions. Animals were intraperitoneally administered PBS (control; n = 3) or IL22Fc (n=4) 24 h prior to infection. The transposon library was cultured in 25 mL of LB liquid media containing 200 μg/mL streptomycin and 50 μg/mL Kanamycin (Strep/Kan) for 3 h at 37 °C with shaking. A higher inoculum size (∼10^8^ CFU) and lower dose of IL22Fc (5 μg/animal) was used in these experiments compared to experiments described in Fig.3 to ensure that there would be sufficient numbers of CFU in intestinal homogenates to allow statistically robust comparisons of the abundances of insertions in the inoculum with the outputs^61^ . Distal SI tissue was homogenized in 1mL of 33% glycerol LB and plated on LB Strep/Kan agar plate and cultured at 37°C overnight. All colonies on those plates were harvested in 2 mL of 33% glycerol LB and stored at −20°C until further processing. Genomic DNA (gDNA) was extracted using gDNA extraction kit (Qiagen), sheared into 400 base pair (bp) using Covaris, and was purified using the DNA clean kit (Qiagen). gDNA fragments were repaired with Blunting enzyme mix (NEB), A-tails were added using Taq polymerase and dATP, and ligated with adaptor sequences. Adaptor-ligated gDNA was amplified using Phusion PCR polymerase with adaptor primers for 30 cycles. After purification, the amplicon was further amplified with var primers and index barcodes. The amplified library with sizes between 300-500 bp were isolated using SPRIselect beads (Beckman-Coulter). Sequencing was performed using a Nextseq1000 with P1 100 cycle cartridge (illumina). Tn-seq data was analyzed using the RTISAn pipeline as previously described^62^.

### Competition assay with barcoded *V. cholerae*

Diluted WT or mutant *V. cholerae* STAMP libraries were cultured on 200 μg/mL streptomycin LB (LB Strep) plates and 4-8 isolated colonies were picked. These single colonies were then individually grown in LB Strep liquid media with shaking at 37 °C overnight; then after adjusting OD600 to 1 with LB media, they were mixed in equal ratios. The barcoded strain mix was stored as a glycerol stock at −80 °C until use.

WT infant mice (Figure 4d) or P4 *Muc2*^+/-^ or *Muc2* ^-/-^ littermate (Figure 6n) were intraperitoneally administered PBS or 5 μg IL22Fc 24 hours before infection. The barcoded strain mix was cultured in LB Strep liquid media with shaking at 37 °C overnight and infant mice were orally inoculated with 1×10^7^ cells of the strain mix and sacrificed at 20 hpi. Homogenates of the distal SI tissue were plated on LB Strep plates, grown at 37 °C overnight, and colonies were collected in 33% glycerol LB. Genomic DNA was extracted from the collected bacteria and DNA barcodes were analyzed according to the protocol described above. To calculate competitive indices, the frequency of each barcode in the output population (from intestinal homogenates) was divided by the frequency in the inoculum. The geometric mean of the resultant ratio of barcode frequencies (output/inoculum) in mutants were compared to those of the WT strain in each individual mouse.

### **Reg3β** bactericidal assay *in vitro* and *in vivo*

An overnight culture of wild type *V. cholerae* was inoculated in LB Strep and incubated at 37°C for 3 hours. The cell density (OD600) was adjusted to 0.1 (∼10^8^ cells/mL) and cells were washed in MES buffer (100mM MES pH5.8, 25mM NaCl) twice. 10^5^ cells were mixed with recombinant Reg3β protein (Sino Biological, 51153-M08H) in MES buffer and incubated at 37°C for 3 h. Cells were resuspended in LB media and either plated on LB Strep agar or growth was monitored in a BioTek Epoc2 Microplate Spectrophotometer (Agilent, CA).

For *in vivo* assays, P5 infant animals were given 33.3ug of Reg3β protein orally 30 min before and 1 hour after mice were orally gavaged with 10^5^ cells of WT *V. cholerae*. Proximal and distal SI tissues were harvested and homogenized in 33% glycerol LB at 16 hours post infection. Serial dilutions of tissue homogenates were plated on LB Strep plates for CFU counting.

### Image analysis

All images were analyzed using Fiji Software version 2.14.0 (https://imagej.net/software/fiji/). Pearson’s correlation coefficient of colocalization between Reg3β and *V. cholerae* were analyzed using Coloc2 plugin (https://github.com/fiji/Colocalisation_Analysis). Z-stack images (0.3μm distance per image) of colocalization between Reg3β and *V. cholerae* were captured with Nikon Microscope Objective Plan Apo VC 100x/1.40 Oil DIC lens and 3D images were built using 3D viewer plugin (https://github.com/fiji/3D_Viewer). *Atoh1*+, *Spdef*+, and EdU+ cells were counted using Find Maxima function with the same setting of Prominence across all compared images.

### Protein extraction, digestion, LC-MS/MS acquisition, and data analysis

The distal SI tissue was collected from control or IL22Fc-treated infant animals and cut open longitudinally. The tissue was incubated in 4 mL of wash buffer [1mM DTT PBS with protease inhibitor (Roche, 4693159001)] on ice for 10 min and vortexed for 20 seconds. After the removal of tissue pieces, intestinal washes were centrifuged at 100 xg for 1 min to remove the large pieces of intestinal contents. The supernatant was concentrated by centrifugation at 4000 xg with 3 kDa MWCO Amicon Ultra centrifugal filter unit for 1 hour. The volume of the concentrate was adjusted to 400 μL with wash buffer. The supernatants were processed for LC-MS/MS analysis using the ENRICH-iST kit (PreOmics, #P.O.00163). Assay details have been described previously^63^, including protein enrichment, denaturation, reduction, alkylation, enzymatic digestion, and peptide cleanup within a single workflow. Briefly, mucus proteins were captured on paramagnetic enrichment beads in binding buffer, followed by multiple bead-based wash steps to remove salts, lipids, and low-molecular-weight contaminants. Bead-captured proteins were denatured, reduced, and alkylated using the supplied lysis buffer at elevated temperature and digested on-bead with a Trypsin/LysC protease mixture. Digestion was terminated by stop solution, and peptides were purified using the solid-phase extraction cartridge with sequential washes to remove impurities. Peptides were eluted in volatile buffer, dried completely by vacuum centrifugation (SpeedVac SPD120), and stored at −80 °C until LC-MS/MS analysis.

Peptides were reconstituted in 5-10 µL of solvent A (2.5% acetonitrile, 0.1% formic acid) and separated by nano-scale reverse-phase liquid chromatography using a custom-packed capillary column (100 µm inner diameter × ∼30 cm length) containing 2.6 µm C18 spherical silica beads and terminated with a flame-drawn electrospray tip^64^. Subsequently, peptides were loaded via an autosampler and eluted using a linear gradient of increasing concentrations of solvent B (97.5% acetonitrile, 0.1% formic acid). Eluting peptides were ionized by electrospray and analyzed on an Orbitrap Velos Pro hybrid ion trap-Orbitrap mass spectrometer (Thermo Fisher Scientific).

MS raw files were searched against the UniProt FASTA databases using Sequest (Thermo Fisher Scientific)^65^. Databases included reversed (decoy) sequences to enable estimation of false discovery rates. Peptide-spectrum matches were filtered to achieve a peptide-level false discovery rate of 1–2%.

Carbamidomethylation of cysteine residues was specified as a fixed modification during database searching. Enzyme specificity was defined as cleavage at the carboxyl terminus of arginine and lysine residues, consistent with digestion by trypsin and LysC. A maximum of ## missed cleavages was allowed. All bioinformatic analyses were performed using Perseus (version 2.0.7.0)^66,67^ as well as standard analytical workflows implemented in R (version 4.5.2).

### Immunoprecipitation and western blot for intestinal contents

The distal SI tissues collected from control and IL22Fc-treated infant animals were cut open longitudinally, were incubated in 4 mL of wash buffer [1mM DTT PBS with protease inhibitor (Roche, 4693159001)] on ice for 10 min, and were vortexed for 20 seconds. After the removal of tissue pieces, intestinal washes were centrifuged at 100 xg for 1 min to remove the large pieces of intestinal contents. The supernatant was concentrated by centrifugation at 4000 xg with 3 kDa MWCO Amicon Ultra centrifugal filter unit for 1 hour. The volume of the concentrate was adjusted to 400 μL with wash buffer and OD600 of wash was measured by SpectraMAX i3x (Molecular Devices). Twenty microliter of intestinal wash was used for immunoprecipitation and western blot. An antibody against Muc2 (ProteinTech, 27675-1-AP) was mixed with Pierce Protein A/G Magnetic Beads (Pierce, 88802) in PBS for 1 hour and further mixed with intestinal wash in 500 μL of IP buffer [0.1% Tween 20 PBS with protease inhibitor] for 3 hours at 4 °C. Beads were washed in 1mL of IP buffer three times, mixed with SDS-PAGE loading buffer containing DTT (NEB, B7703S), and boiled at 95 °C for 5 min. SDS-PAGE (ThermoFisher, WG1403BOX) and transferring to nitrocellulose membrane (ThermoFisher. IB23001) using iBlot 2 (ThermoFisher) were performed according to manufactures’ protocol. After blocking the membrane with 5% skimmilk TBST, the membrane was incubated with antibody against Muc2 (Abcam, EPR23479-47, 1:1000) in TBST at 4 °C overnight. After wash in TBST, the membrane was incubated with IRDye 800CW goat anti-rabbit IgG Secondary Antibody (LICORbio, 926-32211, 1:5000) and UEA-I Fluorescein (vector, FL-1061, 1:5000) at room temperature for 1 hour. After wash in TBST, the fluorescent signal was detected using ChemiDoc MP imaging system (Bio-rad).

### Statistical Analysis

Statistical analysis was performed using Prism10 software version 10.4.0. For the CFU, founding population (Ns), and CFU/Ns comparison in animal infection studies, values were transformed to logarithm 10 before statistical analysis. n represents the number of animals unless otherwise noted in the figure legend.

## Data availability

The single-cell RNA sequence data have been deposited in the NCBI Gene Expression Omnibus (GEO) database under accession number GSE288380. The proteomics data have been deposited in Proteomics Identification database (PRIDE) under project accession number PXD075406 (Project DOI: 10.6019/PXD075406). Extended data tables 1-7 including differentially expressed gene analysis, and single cell pathway analysis (SCPA) are available in Zenodo under accession number 19542157.

## Acknowledgments

We thank members of the Waldor lab for helpful comments on this project and manuscript, Franz Zingl Ph.D. for data in Fig. 5d, Krithika Badarinath, Ph.D. for advice on lineage tracing. We thank Paula Montero Llopis, Ph.D., Praju Vikas Anekal, Ph.D., and Adrienne Wells, Ph.D. in the MicRoN Center at Harvard Medical School for histology, immunohistochemistry and 3D imaging, Ross Tomaino, Ph.D. in the Harvard Taplin Mass Spectrometry Facility, the Harvard Rodent Histology Core for tissue sectioning and staining, the Harvard Biopolymers facility for RNA sequencing, and the Harvard Research Computing Core for providing the computational analysis platform. We are grateful to Anna Velcich at College of Medicine/Albert Einstein Cancer Center for providing us Muc2 knockout mice and to Mikio Hoshino, Ph.D. at National Institute of Neuroscience (PCNP) for Atoh1^Cre(ER-T2)^ mice, and to Genentech for providing us IL22Fc recombinant protein.

## Author contributions

Conceptualization: M.S., Y.H., H. Z., M.K.W.

Methodology: M.S., Y.H., H. Z., Z.L., G.T.

Investigation: M.S., Y.H., Z.L., G.T.

Visualization: M.S., Y.H., Z.L. Funding acquisition: M.K.W. Project administration: M.K.W. Supervision: R.A.S., M.K.W.

Writing – original draft: M.S., Y.H., and M.K.W.

Writing – review & editing: M.S., Y.H., H.Z., Z.L., R.A.S., and M.K.W.

## Competing interests

Authors declare there is no competing interests

### Licence information

This article is subject to the Howard Hughes Medical Institute (HHMI)’s Open Access to Publications policy. HHMI laboratory heads have previously granted a non-exclusive CC BY 4.0 licence to the public and a sublicensable licence to HHMI in their research articles. Pursuant to those licences, the author-accepted manuscript of this article can be made freely available under a CC BY 4.0 licence immediately upon publication. This article is the result of funding in whole or in part by the National Institutes of Health (NIH). It is subject to the NIH Public Access Policy. Through acceptance of this federal funding, the NIH has been given the right to make this article publicly available in PubMed Central upon the Official Date of Publication, as defined by the NIH.

