## Extended data figures for "IL-22 promotes genesis of small intestinal secretory cells that protect against cholera in mice"

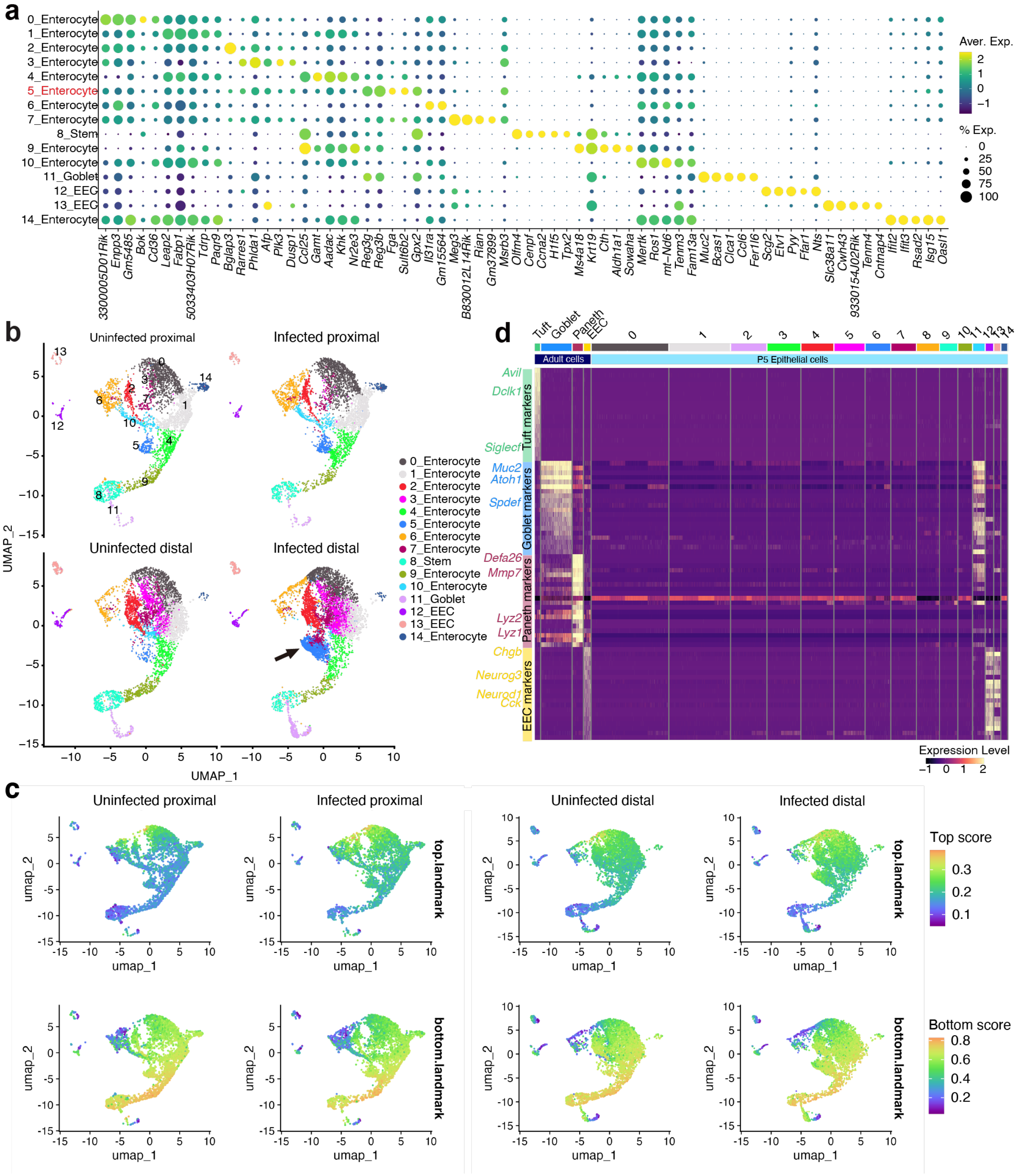

**Extended Data Fig. 1. scRNA-seq profile of the SI epithelial cells in P5 infant**
(**a**) Representative genes for each epithelial cell cluster depicted in Fig.1b. Dot plot indicates the scaled expression levels in color and the percentage of the cells expressing genes in the size of each cluster; EEC (enteroendocrine cells). (**b**) UMAP of the small intestinal epithelial single-cell RNA-seq data of the proximal and distal SI from uninfected and *V. cholerae* infected P5 infant mice (n=3 for each condition, cells are pooled). The arrow points cluster 5 enterocytes in

infected animals in the distal SI. **(c)** Signature scores of top-villus landmark genes (left panels) and bottom-villus landmark genes (right panels) calculated by UCell[69] . Color scales indicate the UCell score. (**d)** Marker genes for tuft cell (green), goblet cell (blue), Paneth cell (red) and enteroendocrine cell (EEC, yellow) and their expression in epithelial cell clusters in adult (Haber et al[68] : GSE92332) and P5 infant mice. Heatmap represents the scaled expression level.

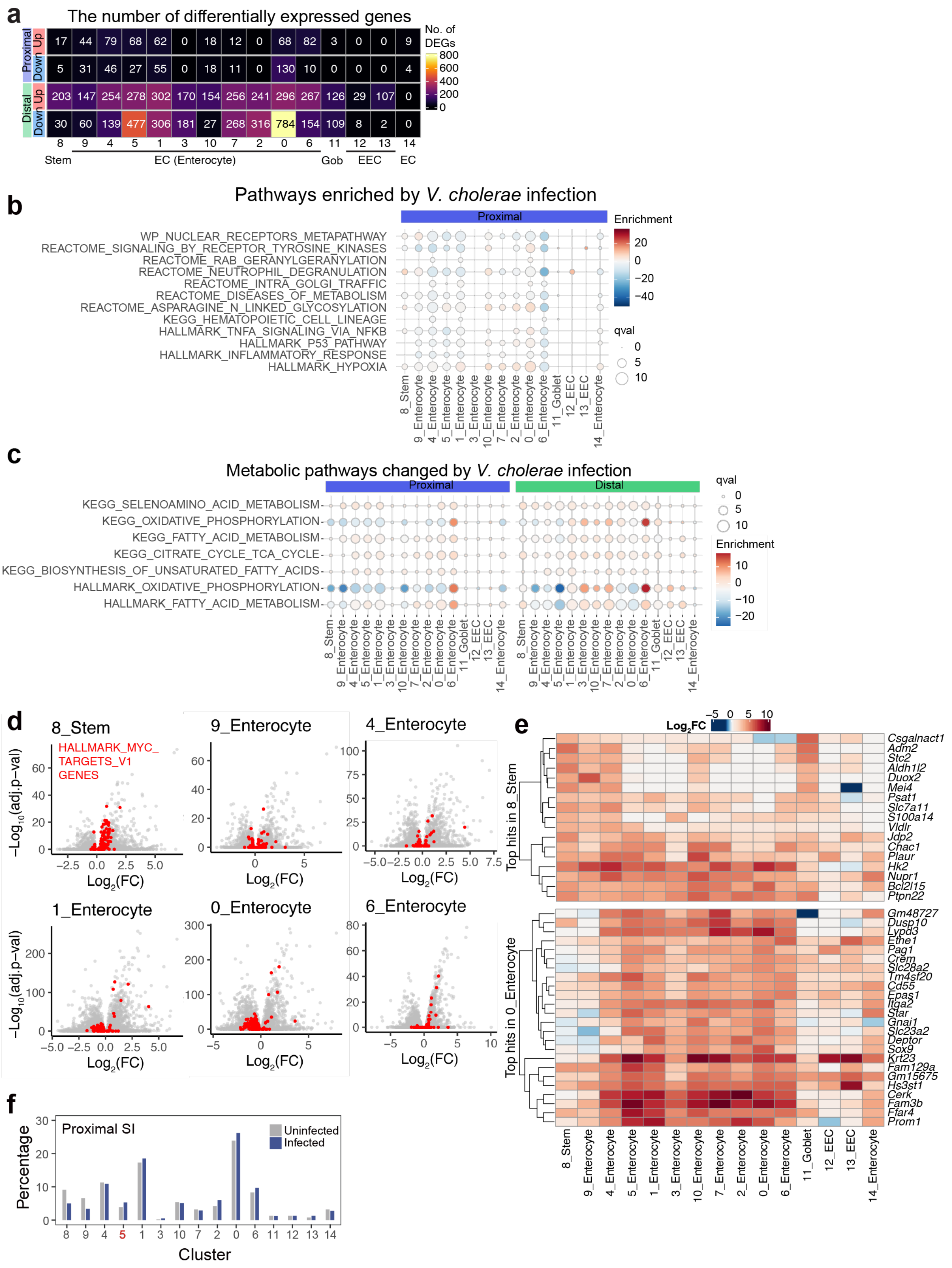


**Extended Data Fig. 2 Epithelial cell-type specific responses to *V. cholerae* infection**

(**a**) Heatmap of the numbers of differentially expressed genes (absolute Fold Change > 1, adjusted p-value < 0.00001, genes on chromosome X and Y were excluded as both, male and female infant mice were used for each condition) in each cluster. Stem, stem cells, EC, enterocytes, Gob, goblet cells, EEC, enteroendocrine cells. Wilcoxon rank-sum test. p-value was adjusted using Bonferroni correction. (**b**) Pathways of the Single Cell Pathway Analysis (Qval > 8.75 in any cluster) in the proximal SI epithelium. Pathways are matched to Fig.1c. Dot plot represents enrichment in color and Qval in circle size. Stem (stem cells); Gob (goblet cells); EEC (enteroendocrine cells). **(c)** HALLMARK and KEGG metabolic pathways altered by *V. cholerae* infection. Dot plot represents enrichment in color and Qval in circle size. (**d**) Volcano plots showing gene expression changes between uninfected and *V. cholerae*-infected conditions. Genes in the HALLMARK_MYC_TARGETS_V1 pathway are highlighted as red. (**e**) Heatmap representing uninfected vs. *V. cholerae*-infected gene expression changes in color. Top: Top-hit DEGs ($Log_2$ fold change > 3 and adjusted p-value < 0.01) in 8_stem, Bottom: Top-hit DEGs in 0_Enterocyte. *Reg3b* and *Reg3g* are excluded. Note that $log_2$ fold change of the genes without any values of differential expression analysis (Seurat's FindMarkers function) are set to zero. (**f**) Percentage of cells in each cluster vs the total cell number in the proximal SI.

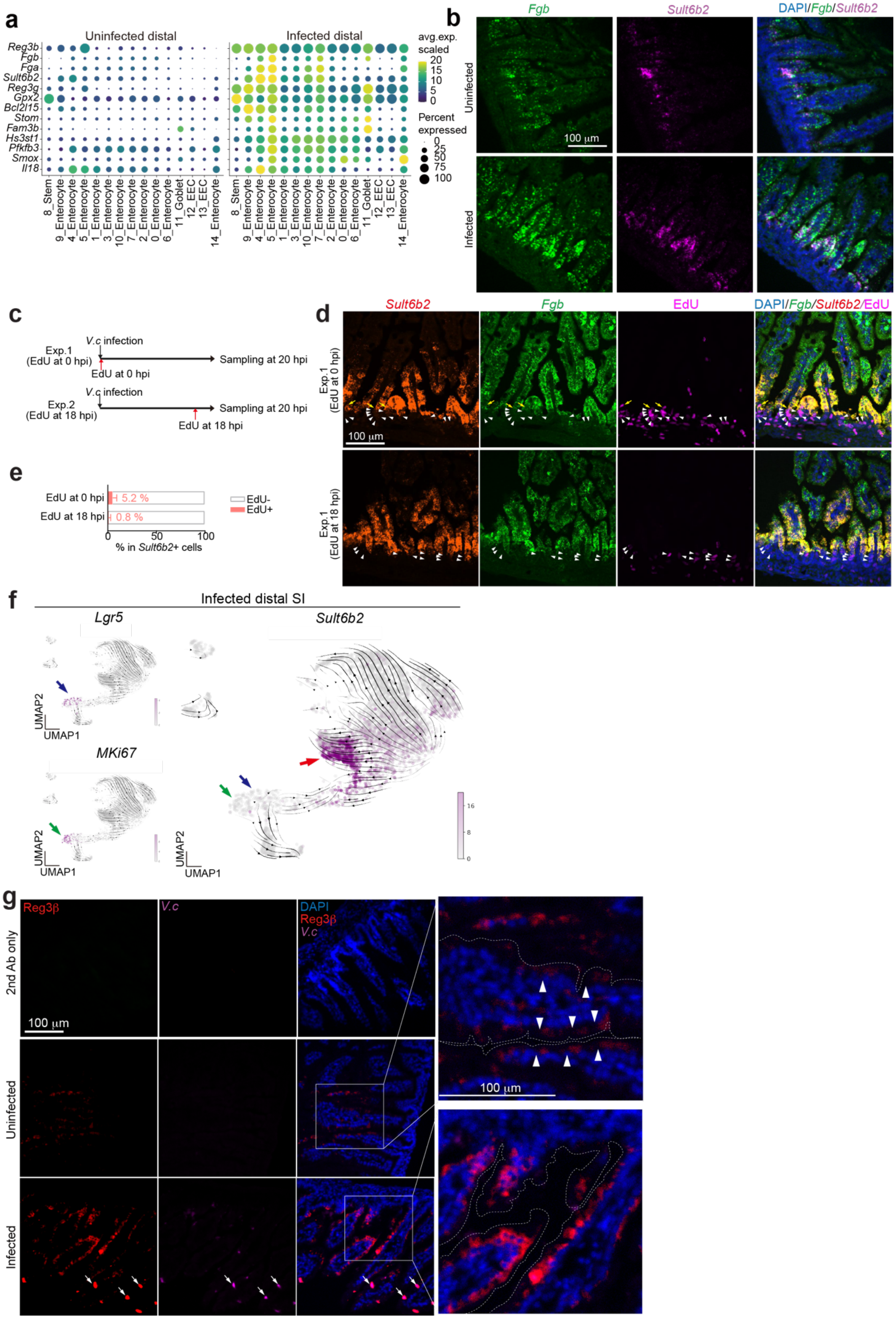
a
Uninfected distal
Infected distal
Reg3b
Fgb
Fga
Sult6b2
Reg3g
Gpx2
Bcl2l15
Stom
Fam3b
Hs3st1
Pfkfb3
Smox
Il18
8_Stem
9_Enterocyte
4_Enterocyte
5_Enterocyte
1_Enterocyte
3_Enterocyte
10_Enterocyte
7_Enterocyte
2_Enterocyte
0_Enterocyte
6_Enterocyte
11_Goblet
12_EEC
13_EEC
14_Enterocyte
avg.exp. scaled
20
15
10
5
0
Percent expressed
0
25
50
75
100
b
Fgb
Sult6b2
DAPI/Fgb/Sult6b2
Uninfected
Infected
100 µm
c
V.c infection
Exp.1 (EdU at 0 hpi)
EdU at 0 hpi
Sampling at 20 hpi
Exp.2 (EdU at 18 hpi)
EdU at 18 hpi
d
Sult6b2
Fgb
EdU
DAPI/Fgb/Sult6b2/EdU
Exp.1 (EdU at 0 hpi)
Exp.1 (EdU at 18 hpi)
100 µm
e
EdU at 0 hpi
EdU at 18 hpi
5.2 %
0.8 %
EdU-
EdU+
0
50
100
% in Sult6b2+ cells
f
Infected distal SI
Lgr5
Sult6b2
MKi67
UMAP2
UMAP1
16
8
0
g
Reg3β
V.c
DAPI
Reg3β
V.c
2nd Ab only
Uninfected
Infected
100 µm

**Extended Data Fig. 3. Identity of cluster 5 epithelial cells**

**(a)** Representative genes for cluster 5 epithelial cells. Dot plots indicate the scaled expression levels in color and the percentage of the cells expressing genes in the size of each cluster of epithelial cells in distal SI in uninfected (left panel) and *V. cholerae* infected animals (right panel).(**b**) Separated images of RNA FISH detecting cluster 5 markers: *Fgb* (green) and *Sult6b2* (magenta). DAPI (blue). Merged images are also shown in Fig. 1g. Shown are representative images of 29 images from 4 uninfected animals and 34 images from 4 infected animals. (**c**) Experimental protocols for experiments with EdU labeling and imaging to investigate the origins of cluster 5 cells. In experiment 1 (Exp. 1), infant mice were given EdU at the same time of infection (0 hpi), and in experiment 2 (Exp 2), infant mice were given EdU at 18 hpi (2 hours before sampling). (**d**) Images of EdU (magenta), co-stained with cluster 5 enterocyte markers, *Sult6b2* (red) and *Fgb* (green), and DAPI. Representative images of 160 images from 4 animals are shown. (**e**) The percentage of EdU+ cells in cluster 5 enterocytes (*Sult6b2*+ cells). 110 (from 4 mice, Exp.1) and 63 images (from 3 mice, Exp.2) were analyzed. Data are presented as mean values ± SD. (**f**) UMAP projection representing the RNA expression of marker genes in the epithelial cell scRNA-seq data from the distal SI in infected animals. RNA-velocity is overlayed on the UMAP. Top left UMAP shows the expression levels of *Lgr5* (stem cells) and the bottom left UMAP shows that of *MKi67* in purple. The right UMAP shows the expression level of cluster 5 enterocyte marker gene *Sult6b2* in purple. Colored arrows on the UMAPs point to *Lgr5*+ cells (blue), *MKi67*+ cells (green), and *Sult6b2*+ cells (red). (**g**) Immunohistochemistry of Reg3β (red), *V. cholerae* (magenta), and DAPI (blue) in the distal SI tissue at 20 hpi. Top row images depict tissues stained with only secondary antibody as a control. Middle and bottom rows are uninfected and *V. cholerae* infected distal SI tissues. White arrows in the images of infected samples show *V. cholerae* microcolonies on the SI villi. Right panels are high magnification images of the merged image. White arrowheads in the high magnification image indicate the expression of Reg3β in the uninfected SI. Dotted line indicates the epithelial cell membrane. Merged images are also shown in Fig. 1k. Representative images of 66 images from 3 animals or 47 images from 3 *V. cholerae*-infected animal are shown. Scales are indicated.

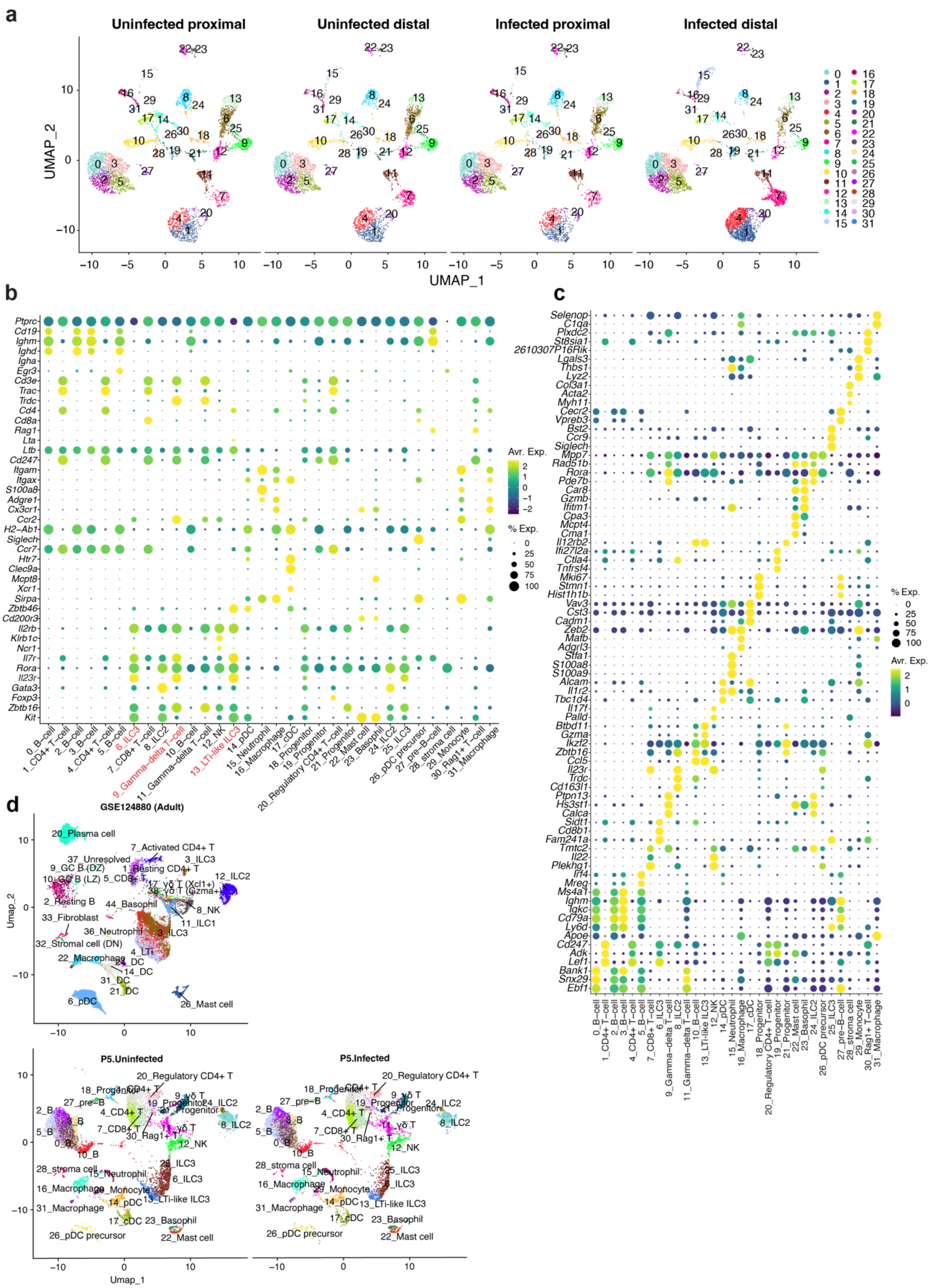
a
Uninfected proximal
Uninfected distal
Infected proximal
Infected distal
UMAP_2
UMAP_1
b
Avr. Exp.
% Exp.
0_B-cell
1_CD4+ T-cell
2_B-cell
3_B-cell
4_CD4+ T-cell
5_B-cell
6_ILC3
7_CD8+ T-cell
8_ILC2
9_Gamma-delta T-cell
10_B-cell
11_Gamma-delta T-cell
12_NK
13_LTi-like ILC3
14_pDC
15_Neutrophil
16_Macrophage
17_cDC
18_Progenitor
19_Progenitor
20_Regulatory CD4+ T-cell
21_Progenitor
22_Mast cell
23_Basophil
24_ILC2
25_ILC3
26_pDC precursor
27_pre-B-cell
28_stroma cell
29_Monocyte
30_Rag1+ T-cell
31_Macrophage
c
d
GSE124880 (Adult)
Umap_2
Umap_1
P5.Uninfected
P5.Infected

**Extended Data Fig. 4. scRNA-seq profile of the SI lamina propria CD45+ cells in P5 infant mice.**

(**a**) UMAP of the CD45+ cells, split by tissue (proximal and distal) and conditions (infected and uninfected). **(b, c)** Canonical marker gene (b) and representative gene expression (c) in each cluster. Canonical marker genes were determined by database search with Tabula Muris[70], PanglaoDB[71] , and literatures[72,73]. Dot plot indicates the scaled expression levels in color and the percentage of the cells expressing genes in the size of each cluster of immune cells in combined data (immune cells in proximal and distal SI from uninfected and infected animals).

(**d**) Integrated UMAP of the immune cells in adult (GSE124880)[17] and infant mice (our data). GSE124880 cell annotations are based on the original data source.

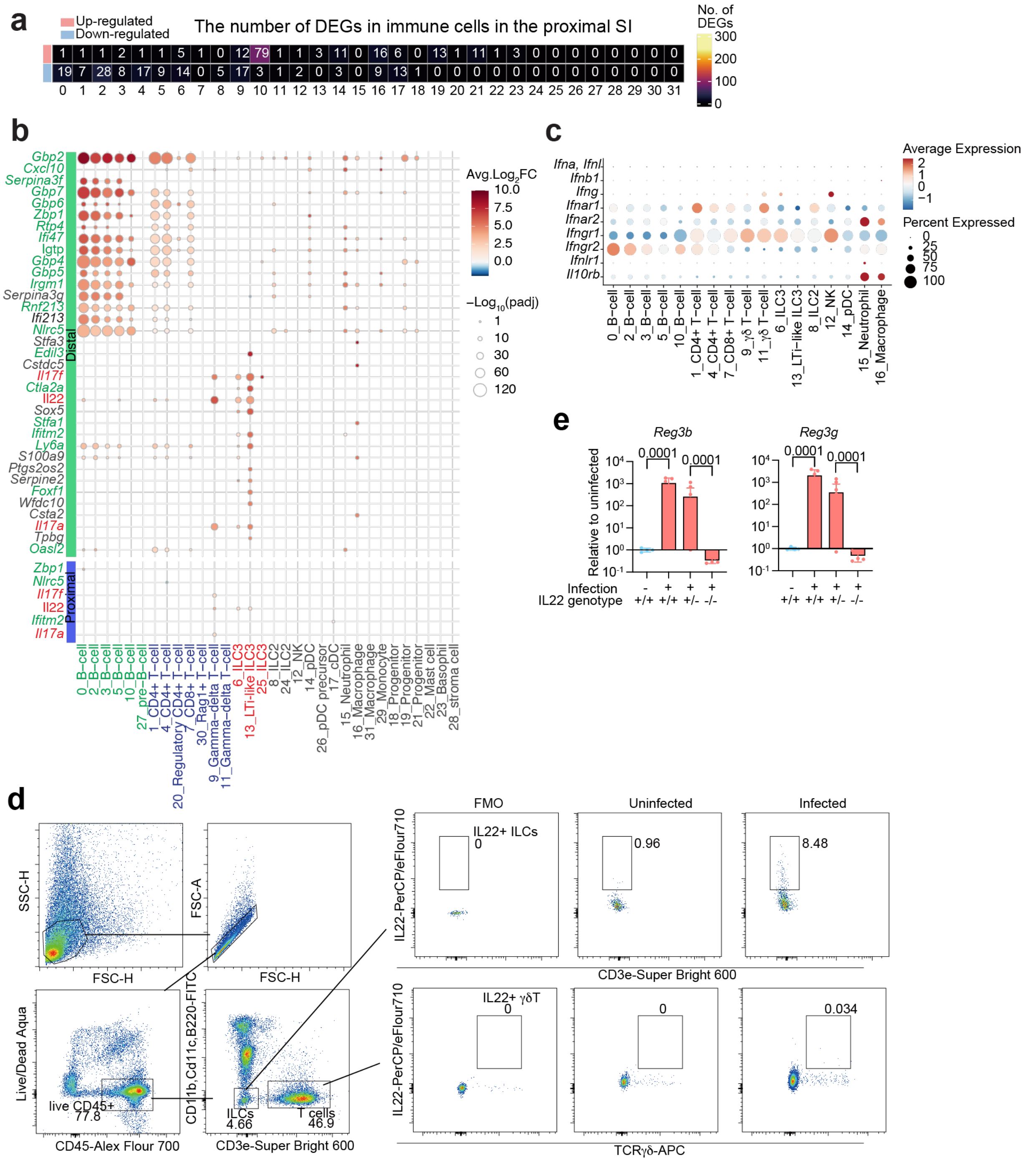

**Extended Data Fig. 5 Cell-type specific transcriptional change**

**(a)** Heatmap of the numbers of differentially expressed genes (absolute Fold Change > 1, adjusted p-value < 0.00001, genes on chromosome X and Y were excluded as both, male and female infant mice were used for each condition, in each cluster). (**b)** Top upregulated genes in the distal small intestine (Fold Change > 3, adjusted p-value < $10^{-20}$, on Autosomes, at least 20% of the cells in cluster express the gene). The expression of the several top-upregulated genes in the distal SI also changes in the proximal SI. *Il22*, *Il17a*, and *Il17f* are highlighted in red. Interferon responsive genes are highlighted in green. Color shows the average $\log_2$ fold change. Dot size represents negative $\log_{10}$ scaled value of adjusted p-value. Wilcoxon rank-sum test. p-value was adjusted using Bonferroni correction (a, b). **(c)** Dot plots showing average expression and percent of expressing cells of interferons and their receptor gene expression in immune cells. Clusters with more than 500 cells (total cell number) are shown. Note that *Ifna* and *Ifnl* were not detected. Color shows the average expression. Dot size represents percent of cells expressing genes in each cluster. **(d)** Representative flow cytometry plots for IL22+ ILCs (CD45+, CD11b-, CD11c-, B220-, CD3e-, IL22+) and γδT cells (CD45+, CD11b-, CD11c-, B220-, CD3e+, TCRγδ+, IL22+) detection in SI lamina propria. FMO: Fluorescence minus one (anti-Il22 antibody) control. **(e)** qPCR analysis of *Reg3b* and *Reg3g* expression in the distal SI from *V. cholerae* infected/uninfected WT C57BL/6 ($IL22^{+/+}$), infected $IL22^{+/-}$ and $IL22^{-/-}$ littermate animals. Expression levels were normalized by β-Actin expression and shown as relative expression to uninfected $IL22^{+/+}$ group. Biological replicates: uninfected $IL22^{+/+}$ (n = 4); infected $IL22^{+/+}$ (n = 5); infected $IL22^{+/-}$ (n = 10); infected $IL22^{-/-}$ (n = 5). Data are presented as mean values ± SD. Two-sided one-way ANOVA with Holm-Šídák's multiple comparisons test.

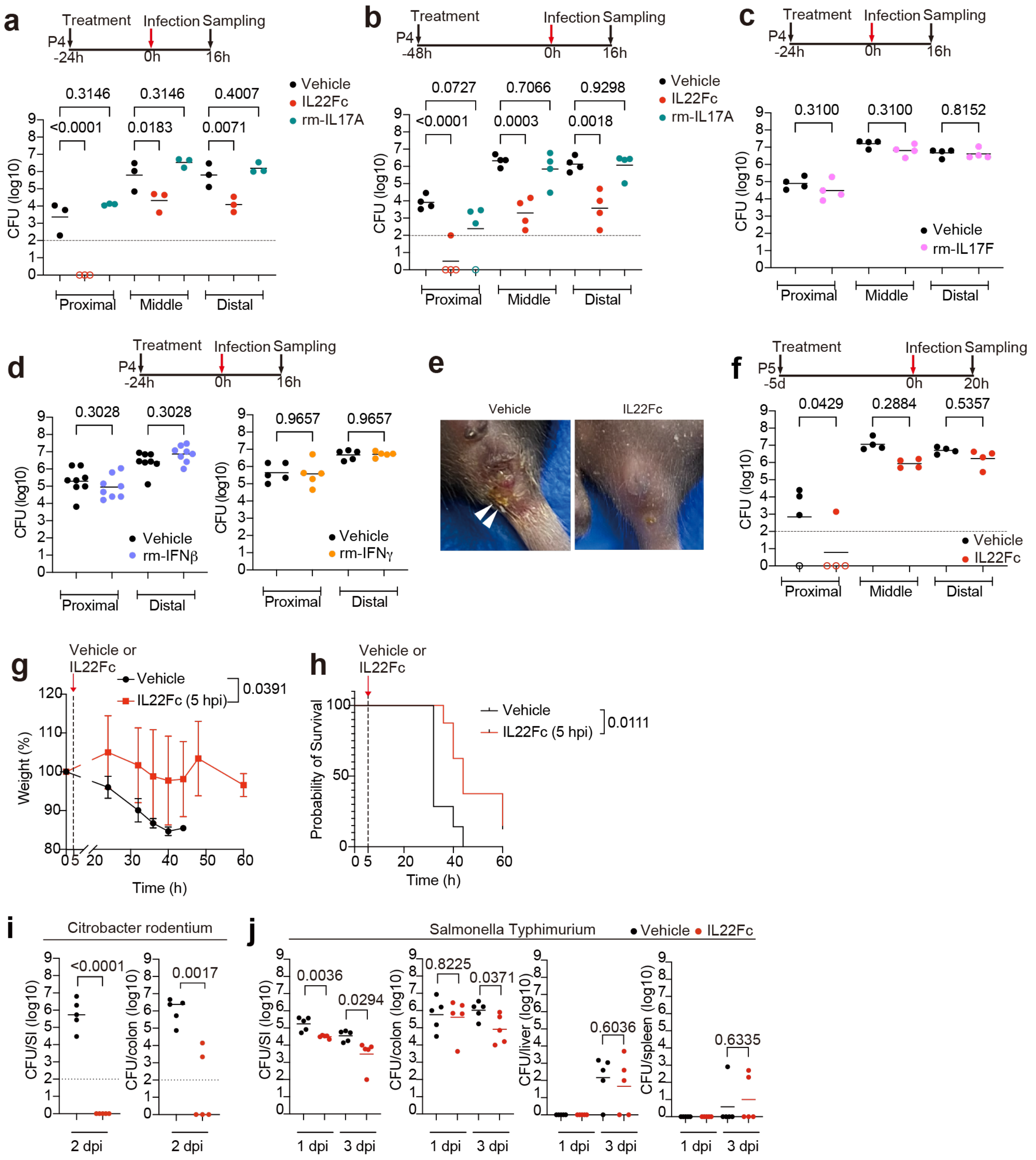

**Extended Data Fig. 6. Dosage regimen-dependent efficacy of IL22Fc on *V. cholerae* colonization and effect of IL22Fc on colonization with other enteric pathogens**

**(a, b)** *V. cholerae* burden in the SI tissues in animals treated with vehicle (black), 5µg IL22Fc (red), or 5 µg mouse recombinant IL17A (rm-IL17A) (green) 24 h (n = 3/group) (a) or 48 h (n = 4/group) (b) before infection. **(c, d)** *V. cholerae* burden in the SI tissues in animals treated with vehicle (black) or 5µg mouse recombinant IL17F (rm-IL17F) (pink) (n = 4/group) (c), 5000 unit of recombinant mouse IFNβ (rm-IFNβ, purple, left panel) (n = 8/group) or 2 µg of recombinant mouse IFNγ (rm-IFNγ, orange, right panel) (n = 5/group) (d) 24 h before infection. **(e)** Diarrhea in *V. cholerae*-infected infant mice at 48 hpi. White arrowheads in vehicle-treated animals point to clear diarrheal fluid on the anus. **(f)** *V. cholerae* burden in the SI tissues in animals treated with vehicle (black) or 5μg IL22Fc (red) 5 days before infection (n = 4/group). **(g, h)** Percent of body weight compared to that at 0 hpi (g) and survival (h) after *V. cholerae* infection. Mice were infected with WT *V. cholerae* and administered vehicle or IL22Fc i.p. at 5 hpi. Data are represented as mean ± SD for biological replicate (Vehicle: n = 7; IL22Fc: n = 8), Two-way ANOVA (g). Log-rank test (h). **(i)** *C. rodentium* burden in the entire SI and colon in vehicle (black) or IL22Fc-treated (red) infant animals at 2-days post infection (dpi). Bars indicate the geometric mean for biological replicates (n = 5/group), Unpaired t-test. The dotted lines indicate the detection limit. **(j)** *S.* Typhimurium burden in the entire SI, colon, liver, and spleen in vehicle (black) or IL22Fc-treated (red) infant animals at 1- or 3 dpi. One-way ANOVA. (a, b, c, d, f, j) Bars indicate the geometric mean for biological replicates. p-values are shown.

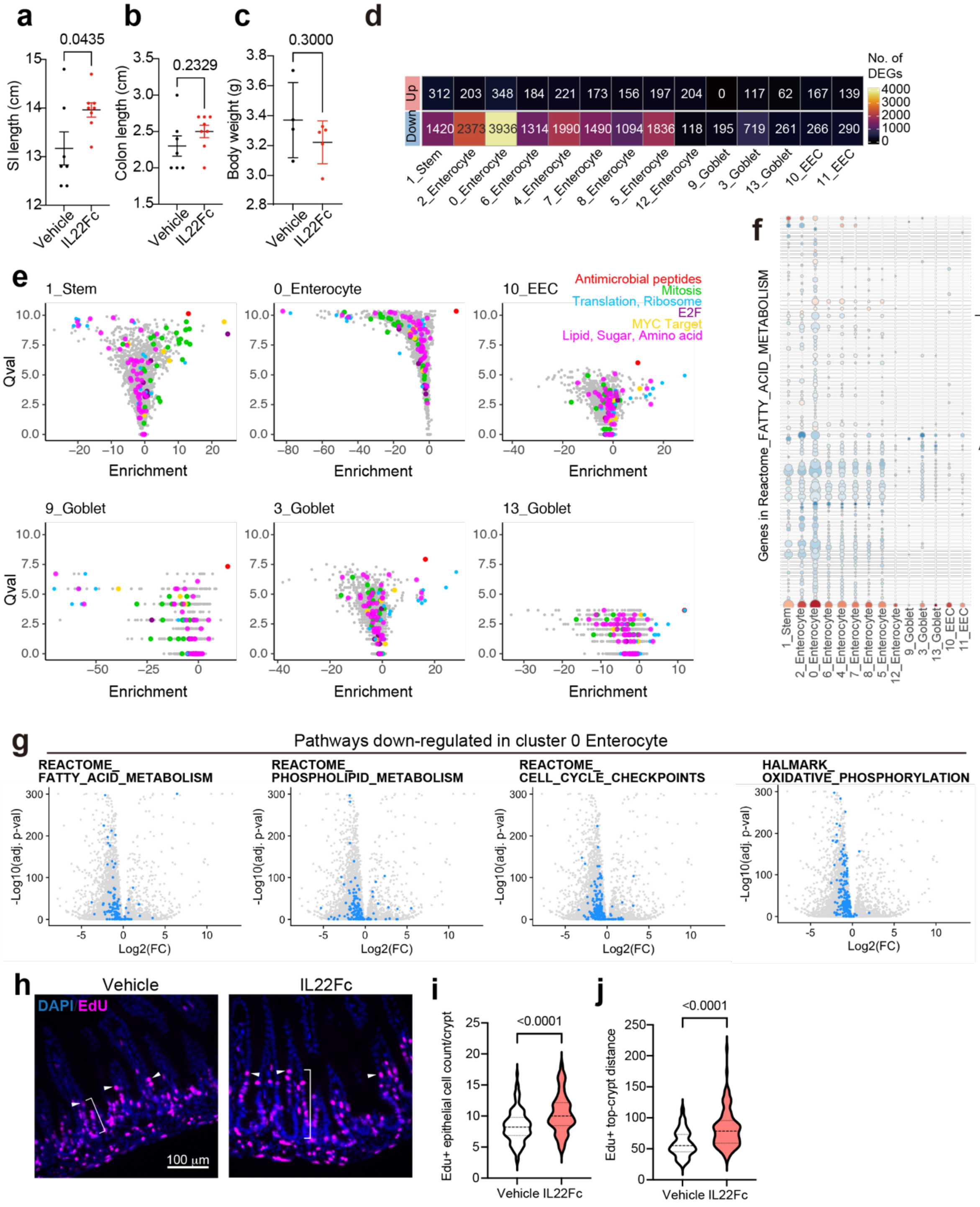


**Extended Data Fig. 7. IL22Fc administration alters the SI epithelium**

(**a-c**) SI (a) and colon (b) length, and body weight (c) of animals 40 hours after vehicle (black) or IL22Fc (red) administration. Data are represented as mean ± SD for biological replicate (Vehicle: n = 7, IL22Fc: n = 8) for the measurement of SI and colon length, and (Vehicle: n = 4; IL22Fc: n = 5) for the measurement of body weight. Two-sided unpaired t-test. p-value is indicated in individual plots. (**d**) Heatmap of the numbers of differentially expressed genes (absolute Fold Change > 1, adjusted p-value < 0.00001, genes on chrX and Y were excluded as male and female infants were mixed) in each cluster. **(e)** Volcano plots showing fold change and q-value of Single Cell Pathway Analysis results. Colors represent pathways: Antimicrobial peptides (red), Mitosis (yellow green), Translation and Ribosome (light blue), E2F (purple), MYC Target (yellow), and Lipid, Sugar, Amino acid (pink). (**f)** Dot plot represents control vs. IL22Fc-treated gene expression changes in the REACTOME_FATTY_ACID_METABOLISM pathway. Average of $\log_2$ fold change in IL22Fc-treated compared to control is shown in color and negative $\log_{10}$-scaled adjusted p-value is shown in the size of each cluster. (**g**) Volcano plots show differentially expressed genes between IL22Fc- vs. control-treated animals in cluster 0 enterocytes. The x- and y-axis indicate $\log_2$ fold change compared to control animals and negative $\log_{10}$ adjusted p-value, respectively. Genes in REACTOME_FATTY_ACID_METABOLISM, REACTOME_PHOSPHOLIPID_METABOLISM, REACTOME_CELL_CYCLE_CHECKPOINTS, and HALLMARK_OXIDATIVE_PHOSPHORYLATION pathways are highlighted in blue. Wilcoxon rank-sum test with Bonferroni correction (d, f, g). (**h**) Representative images of 5-ethynyl 2´-deoxyuridine (EdU) positive cells in the distal small intestine. P4 infant mice were intraperitoneally treated with vehicle or IL22Fc, and EdU. Mice were sacrificed at 36 h post administration. DNA-incorporated EdU was stained with Alexa 647 (magenta). White arrowheads point to the top EdU-positive cells along the crypt-villus axis. The white bracket shows the distance from the bottom of the crypt to the top of the EdU-positive cells. Scale bar: 100 μm. **(i)** Enumeration of EdU positive cells per crypt in the control vs. IL22Fc-treated animals. Vehicle: n = 117 images from 4 animals, IL22Fc: n = 118 images from 4 animals. Two-sided unpaired t-test. **(j)** Measurement of the distance from the bottom of the crypt to the top EdU-positive cells. The migration velocities are coupled to cell proliferation rates[74]. Vehicle: n = 120 images from 4 animals, IL22Fc: n = 120 images from 4 animals. Two-sided unpaired t-test.

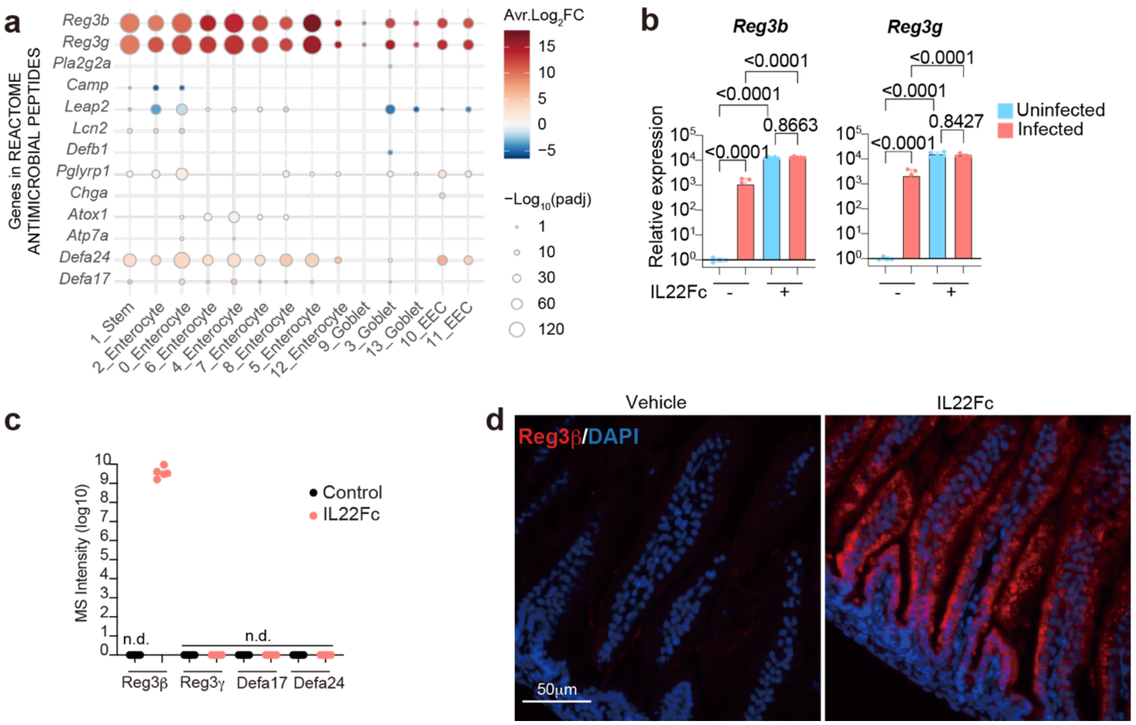


**Extended Data Fig. 8. IL22Fc administration increased the expression of Reg3β in the epithelial cells**

**(a)** Relative expression of the DEGs in the Reactome antimicrobial peptide pathway. Color code and the scale of the dot size indicate average $\log_2$ fold change and negative $\log_{10}$ scaled adjusted p-value in IL22Fc-treated animals, respectively. Wilcoxon rank-sum test with Bonferroni correction. (**b**) qPCR analysis of *Reg3b* and *Reg3g* in the distal SI tissue. Data is normalized by the expression of β-Actin and presented as relative to IL22Fc-untreated/uninfected control. Biological replicate: uninfected IL22Fc(-) (n = 4); infected IL22Fc(-) (n = 5); uninfected IL22Fc(+) (n = 5); infected IL22Fc(+) (n = 5). Two-sided one-way ANOVA. Holm-Šídák's multiple comparisons test.. mean ± SD. (**c**) $\text{Log}_{10}$-scaled mass-spectrometry signal intensity of Reg3β, Reg3γ, Defa17 and Defa24 peptides in intestinal wash proteomics analysis in control- (black) vs IL22Fc-treated animals (red). n.d.: not detected. **(d)** Representative images of immunohistochemistry of Reg3β protein (red) and DAPI (blue) in the small intestine from control or IL22Fc-treated animal. Shown are representative images of 54 images from 4 vehicle-treated animals and 65 images from 4 IL22Fc-treated animals. Scale bar: 50 μm.

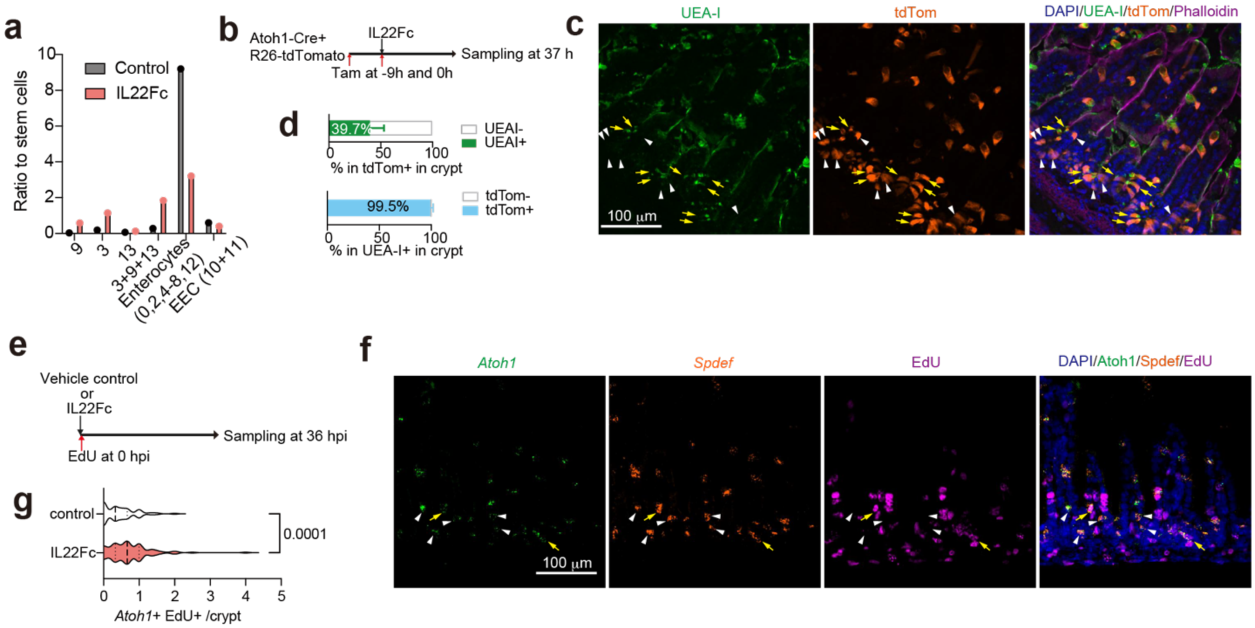


**Extended Data Fig. 9 IL22Fc promotes secretory progenitor cells proliferation**
(**a**) Ratio between indicated cell cluster vs stem cell clusters in scRNA-seq data in Figure 5a. Cell counts per cluster was divided by the count of stem cells. (**b**) Experimental protocol for testing if IL22Fc-induced crypt associated goblet cell arise from the *Atoh1*+ secretory lineage in Atoh1Cre(ER-T2);Rosa26R-LSL-TdTomato animals. Tamoxifen was administered -9 and 0 post IL22Fc administration. Tissue was sampled 37 h post IL22Fc administration. (**c**) Tissue sections were stained with UEA-I as a marker for mucus+ cells (green), and DAPI (blue). White arrowheads and yellow arrows indicate tdTomato+ UEA-I- cells and tdTomato+ UEA-I+ cells, respectively. Shown are representative images of 92 images from 3 animals. (**d**) The percentage of UEA-I+ cells in tdTomato+ cells (top), and tdTomato+ cells in UEA-I+ cells (bottom). 92 images of the small intestine in Atoh1$^{Cre(ER\text{-}T2)}$;Rosa26R$^{LSL\text{-}tdTomato}$ mice (n = 3) were analyzed. Data are presented as mean values ± SD. (**e**) Experimental protocol for testing if IL22Fc treatment promotes proliferation of crypt-associated *Atoh1*+ cells; EdU was administered at 0 hpi. (f) Tissue was stained with RNA FISH probes for the secretory lineage markers *Atoh1* and *Spdef* and EdU was detected with click chemistry. White arrowheads and yellow arrows indicate *Atoh1*+ EdU- cells and *Atoh1*+ EdU+ cells, respectively. Shown are representative images of 100 images from 4 IL22Fc-treated animals. (**g**) Enumeration of *Atoh1*+ EdU+ cells per crypt. 140 and 100 images of the small intestine in control (n = 4) vs IL22Fc-treated animals (n = 4) were analyzed. Two-sided Welch's t-test. p-value is shown.

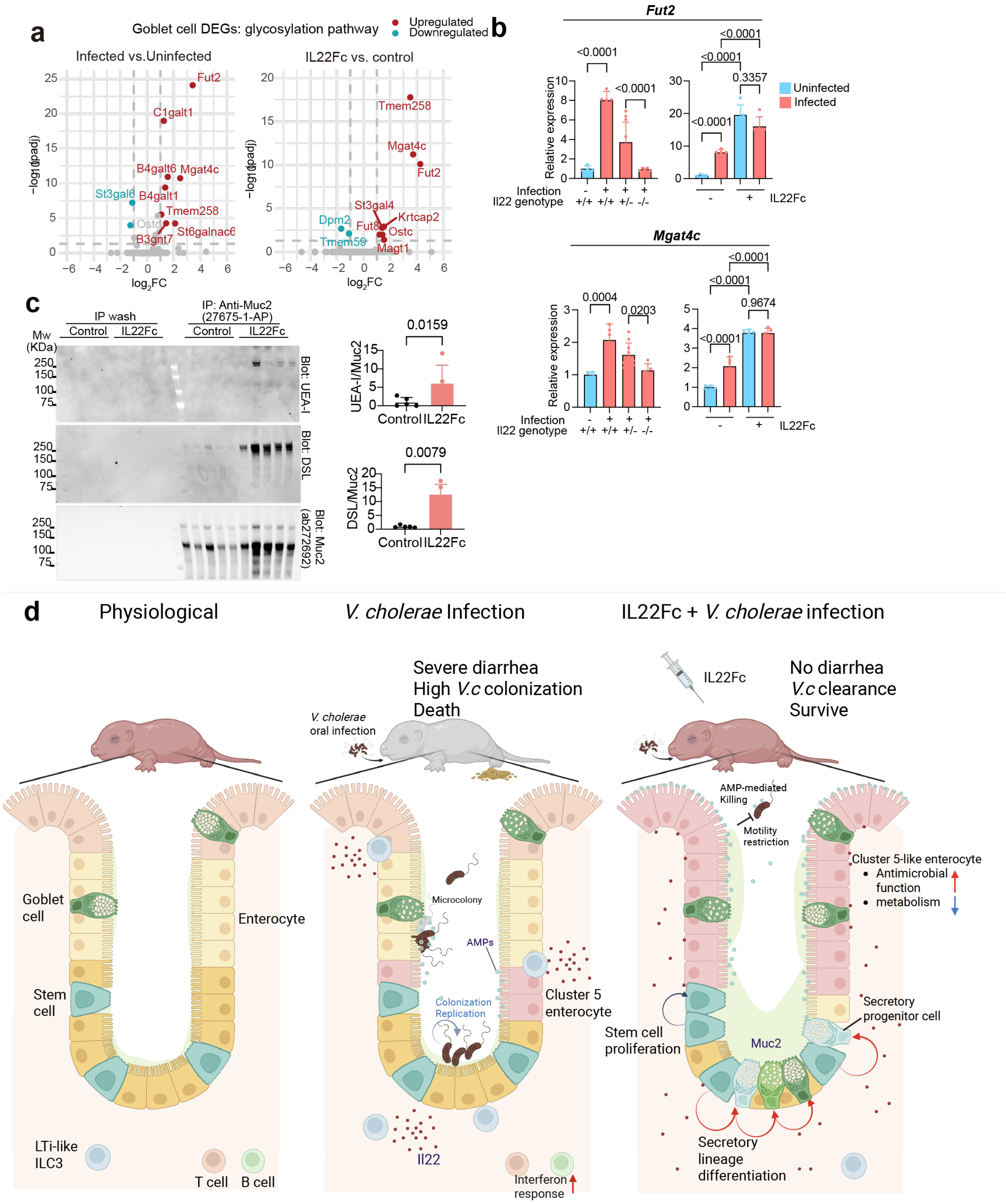


**Extended Data Fig. 10. IL22Fc administration modifies mucus and protects against experimental cholerae**

(**a**) Volcano plots represent differentially expressed genes associated with glycosylation (GO:0006486) in goblet cells in uninfected vs *V. cholerae* infected (left), or control vs IL22Fc-treated animals in the scRNA-seq data (right). Wilcoxon rank-sum test with Bonferroni correction. (**b**) Expression of glycosylation genes in the distal SI tissues in uninfected (n = 4) or *V. cholerae*-infected WT C57BL/6 (+/+) (n = 5), IL22 heterozygous (+/-) (n = 10), or IL22 knockout (-/-) (n = 5) animals (left), and uninfected IL22Fc (-) (n = 4), infected IL22Fc (-) (n = 5), uninfected IL22Fc (+) (n = 5), vs infected IL22Fc (+) (n = 5) animals (right). Data are represented as mean ± SD. One-way ANOVA. (**c**) Western blot image of immunoprecipitation of Muc2 and its modification with fucose (UEA-I) and *N*-acetylglucosamine (DSL) in the SI luminal wash. Relative abundance of modification (UEA-I/Muc2, DSL/Muc2) in Muc2 (250kDa band) is quantified using densitometry and shown as relative to control (right graph) (n = 5/group). Two-sided Mann-Whitney test. Data are represented as mean ± SD in bar graphs. p-values are shown. (**d**) The intestinal response to *V. cholerae* infection and the mechanism of IL22Fc-mediated protection. Infant mice lack Paneth cells but almost all major immune cell types are present except for plasma cells (left). *V. cholerae* infection recruits LTi-like ILC3 to the lamina propria and epithelial layer, and activates IL22 secretion. *V. cholerae* infection elevates the abundance of a subset of epithelial cells (cluster 5) specialized in defense responses such as production of Reg3β, which binds *V. cholerae* microcolonies and kills *V. cholerae.* Despite these defense responses, infant mice die from cholera-like diarrhea (middle). IL22Fc administration induces nearly all enterocyte subsets to express defense associated phenotypes and antimicrobial function is up-regulated and lipid metabolism is down-regulated. IL22Fc also promotes differentiation of stem cells into the secretory lineage cell increasing production of Muc2 mucin. Muc2 restricts *V. cholerae* motility and reduces pathogen association with the epithelium. These mechanisms impair *V. cholerae* colonization, preventing diarrhea and death. (Fig. d is created with BioRender.com.)

### Supplementary information

**Supplementary Table 1. Excel file containing marker genes for clustering in epithelial and immune cell scRNA-seq.**

**Supplementary Table 2. An Excel file containing differentially expressed genes in epithelial cells in the proximal SI comparing uninfected vs *V. cholerae* infected animals.**

**Supplementary Table 3. An Excel file containing differentially expressed genes in epithelial cells in the distal SI comparing uninfected vs *V. cholerae* infected animals.**

**Supplementary Table 4. An Excel file containing differentially expressed genes in immune cells in the proximal SI lamina propria comparing uninfected vs *V. cholerae* infected animals.**

**Supplementary Table 5. An Excel file containing differentially expressed genes in immune cells in the distal SI lamina propria comparing uninfected vs *V. cholerae* infected animals.**

**Supplementary Table 6. An Excel file containing the in vivo Tn-seq analysis result.**

**Supplementary Table 7. An Excel file containing differentially expressed genes in epithelial cells in the distal SI lamina propria comparing IL22Fc-treated vs vehicle-treated animals.**

**Supplementary Table 8. An excel file containing luminal wash proteomics data.**

**Supplementary Table 9. An Excel file containing oligonucleotide sequences for the construction of deletion mutant strains of *V. cholerae* and primer sequences for qPCR**
